# A Protocol for Single Nucleus RNAseq from Frozen Skeletal Muscle

**DOI:** 10.1101/2022.10.02.510482

**Authors:** Tyler GB Soule, Carly S Pontifex, Nicole Rosin, Matthew M Joel, Sukyoung Lee, Minh Dang Nguyen, Sameer Chhibber, Gerald Pfeffer

## Abstract

Single cell technologies are a method of choice to obtain vast amounts of cell-specific transcriptional information under physiological and diseased states. Myogenic cells are resistant to single nucleus RNA sequencing (snRNAseq) due to their large, multinucleated nature. Here, we report a novel, reliable, and cost-effective method to analyze frozen human skeletal muscle by snRNAseq. This method yields all expected cell types for human skeletal muscle and works on tissue frozen for long periods of time and with significant pathological changes. Our method is ideal for studying banked samples with the intention of studying human muscle disease.

## 1. Introduction

Skeletal muscle is a complex tissue composed of around 40% of an adult human’s mass. It contains many cell types, including satellite cells, myotubes, myogenic precursors, fibroadipogenic progenitors (FAPs), fibroblasts, endothelial cells (ECs), pericytes, adipocytes, immune cells, and smooth muscle. Single cell sequencing technologies are becoming an important tool to interrogate the individual transcriptomes of cell types in otherwise complex heterogenous tissues. Increasingly, this technique is proving well suited to simultaneously examining many cell types, uncover rare disease states, and identify subpopulations with distinct functions (Birnbaum, 2018). Few studies have examined human skeletal muscle at the single cell level (Barruet et al., 2020; De Micheli, Spector, Elemento, & Cosgrove, 2020; Orchard et al., 2021; Rubenstein et al., 2020), however, its utility is clear for basic and medical science studies of muscle in health and disease.

Many challenges exist to developing a method to profile all cell types in human skeletal muscle. Since muscle cells are one of the few multinucleated cells in the human body, there are major limitations to using single cell RNA sequencing (scRNAseq) on muscle. The most commonly used scRNAseq method relies on individual cells being encapsulated in a droplet during processing. Due to microfluidic size constraints of 10X and FACS machines, only cells smaller than 70 µm can be sorted. Multinucleated muscle cells are exceedingly large and it is not possible to FACS sort them or pass them through filters or microfluidics (Zeng et al., 2016). For this reason, single cell data typically include very low proportions of myofibers (De Micheli et al., 2020). Conversely, isolating nuclei from muscle yields very high proportions of myonuclei (Dos Santos et al., 2020; Orchard et al., 2021; Petrany et al., 2020). Single nuclear RNA sequencing is the best option if the experimental objective is to obtain sequencing data from all cell types in a muscle biopsy.

Single nuclear RNA sequencing (snRNAseq) has been demonstrated by multiple groups to yield comparable data to scRNAseq (Slyper et al., 2020; Zeng et al., 2016). snRNAseq recovers the same cell types present in a tissue, but in different proportions (Slyper et al., 2020). A notable difference is that nuclei are enriched for non-coding RNAs like lncRNA and miRNA (Zeng et al., 2016). snRNAseq can also target up to 40,000 nuclei, enabling analysis of many nuclei at once (Orchard et al., 2021).

Maintaining the integrity of the nuclei and the RNA contained within is critical if an experiment is to yield useful data. For this reason, using freshly isolated tissue is the preferred approach for scRNAseq and snRNAseq. Obtaining fresh human muscle samples is possible but not always practical. Muscle biopsy is an invasive procedure that is usually performed for clinical and diagnostic purposes. Obtaining the fresh sample requires advance consent and extensive coordination with health providers for a highly time-sensitive protocol. Using frozen tissue negates the requirement to process biopsies as soon as they are obtained. It removes time-sensitivity for obtaining consent and sample handling, which can both be managed after the sample has been collected and frozen. This makes banked tissues available for snRNAseq. The use of frozen tissues also optimises resource utilisation, since the decision to pursue snRNAseq may be made after additional information is available from tissue histopathologic or other molecular studies.

Few papers have applied snRNAseq to skeletal muscle (Dos Santos et al., 2020; Jiang et al., 2020; Kim et al., 2020; Orchard et al., 2021; Petrany et al., 2020; Zeng et al., 2016). Of these, three groups isolated nuclei directly from mouse muscle (Dos Santos et al., 2020; Kim et al., 2020; Petrany et al., 2020), two used cell cultures (Jiang et al., 2020; Zeng et al., 2016), and only one used human skeletal muscle, although they focussed on analyzing accessible chromatin regions by means of ATAC-seq (Orchard et al., 2021). To isolate nuclei directly from muscle, these reports used fiber dissection and dounce homogenization (Dos Santos et al., 2020), or a mix of various homogenization methods and detergents (Orchard et al., 2021; Petrany et al., 2020).

Here we report a simple, fast, and reliable method to isolate over 7000 nuclei on average, an excellent yield compared to other methods (Dos Santos et al., 2020; Kim et al., 2020; Orchard et al., 2021; Petrany et al., 2020), and isolate nuclei from muscle that has been frozen for up to 15 years. Our protocol uses mechanical disruption, filtration, and fluorescence assisted nuclei sorting (FANS) to preserve nuclear membrane and RNA integrity. The method can be applied to different human muscle types in healthy and disease states.

## 2. Materials and Methods

### Human Skeletal Muscle Sample Collection

Research ethics board approval was obtained for this study from the University of Calgary Conjoint Health Research Ethics Board (REB16-2196). Additional muscle samples for research were collected during clinical muscle biopsy procedures, with advance written informed consent. We also used banked muscle tissues from prior procedures, again with written informed consent (REB15-2763). Samples were stored at -80°C after devitalisation. We included 2 samples from participants with normal histopathology and 4 samples from participants with myopathic abnormalities (Joyce, Oskarsson, & Jin, 2012) (m1 having minor abnormalities and M1, M2, and M3 having more severe abnormalities, **Table 1**).

**Table 1.** Summary of Human Samples: Chosen samples represent both sexes, a range of ages, and various disease states of muscle.

| Sample | Disease State | Sex | Age at biopsy | Muscle |
| --- | --- | --- | --- | --- |
| **C1** | Healthy Control | M | 48 | Deltoid |
| **C2** | Healthy Control | F | 33 | Vastus lateralis |
| **m1** | Mild myopathic changes | M | 64 | Deltoid |
| **M1** | Major myopathic changes | F | 52 | Vastus lateralis |
| **M2** | Major myopathic changes | M | 59 | Deltoid |
| **M3** | Major myopathic changes | M | 36 | Deltoid |

### Materials

The requisite materials and reagents used for this protocol are listed (along with suppliers and product numbers) in **Table 2**.

**Table 2.** Summary of materials and reagents with suppliers and product numbers.

| Reagent or Resource | Source | Identifier |
| --- | --- | --- |
| **Consumables** |  |  |
| Polystyrene Petri Dishes | VWR | 25384-088 |
| Stainless Steel Disposable Scalpels | Integra | 4-420 |
| 2 mL 1.44 mm ceramic bead tubes | Bertin | P000912-LYSK0-A |
| 40 µM FlowMi Cell Strainers | SP Bel-Art | 136800040 |
| RNase OUT Ribonuclease Inhibitor | Invitrogen | 10777-019 |
| 5 mL round-bottom tube | Falcon | REF 352058 or REF 352063 |
| DAPI (10 µg/mL) | Sigma Aldrich | D9542 |
| RNase Away |  |  |
| 1000 µL Wide-Bore Pipet Tips | Sigma Aldrich | AXYT1005WBC |
| Single Cell Commercial Assay |  |  |
| Chromium Next GEM Chip G Single Cell Kit, 16 rxns | 10X Genomics | 1000127 |
| Dual Index Kit TT Set A, 96rxn | 10X Genomics | 1000215 |
| Chromium Next GEM Single Cell 3'Kit v3.1, 4 rxns | 10X Genomics | 1000269 |
| Library Construction Kit, 4 rxns-1000196 | 10X Genomics | 1000196 |
| Tube, Dynabeads MyOne SILANE | 10X Genomics | 2000048 |
| 10X Chromium Controller | 10X Genomics | 1000202 |
| **Equipment** |  |  |
| Minilys Homogenizer | Bertin |  |
| FACSaria Fusion | BD |  |
| ST 40R Centrifuge | Sorvall |  |
| TX-750 Rotor | Thermo Scientific |  |
| **Buffers** |  |  |
| Nuclei EZ Lysis Buffer with RNase Inhibitor | Invitrogen | 10777019 |
| Nuclei Lysis Buffer |  |  |
| 0.1X Nuclei EZ Lysis buffer in 1X PBS with 0.2 U/µL RNase Inhibitor |  |  |
| Nuclei Wash and Resuspension Buffer |  |  |
| 1X PBS, 1.5% BSA, 0.2 U/µL RNase Inhibitor, 10 µg/mL DAPI |  |  |
| **Software and Algorithms** |  |  |
| CellRanger | 10X Genomics | Version 6.1.2 |
| STAR | (Dobin et al., 2013) | Version 2.7.2a |
| R | R Core | Version 4.1.2 |
| Rstudio | R Core | Version 1.4.1106 |
| Seurat | (Hao et al., 2021) | Version 4.1.0 |

### Preparation

- 1 Set a swinging bucket rotor to 4°C (fixed bucket rotors may result in loss of nuclei to the sides of the tube rather than collecting the nuclei as a pellet at the base of the tube)
- Prepare Nuclei Lysis Buffer (see Materials) at a minimum of 750 µl per sample
- Prepare Nuclei Wash and Resuspension buffer (see Materials) at a minimum of 550 µl per sample
- Prepare Nuclei Wash and Resuspension buffer (see step 14) for sorting at a minimum of 100 µl per sample

### Tissue Homogenization

Note: Maintain samples on ice wherever possible. All steps should be performed using a 1 mL pipette with a cut tip to create a wider bore unless otherwise indicated. Wide bore tips may be purchased (see materials list) or generated by cutting the tip of a standard 1000ul pipette tip to make a bore approximately 1.3 mm in diameter.

1. On a petri dish in a 4°C room, cut approximately 60 mg of muscle tissue and mince with a sterile scalpel until muscle is reduced to a slurry
2. Transfer muscle to a 2 mL bead tube and add 500 µL Nuclei Lysis buffer (see Materials)
3. Lyse with mechanical homogenizer at 3000 rpm for 2×5 seconds with a 5 second break in between cycles on ice
4. Pipet off the supernatant into a Lo-Bind 1.5 mL Eppendorf. Ensure to minimize residual tissue fragments transferred with the supernatant as this will interfere with filtration.
5. Add 250 µL Nuclei Lysis buffer to the tube containing residual tissue and lyse again for 10s at 3000 rpm. Repeat step 4.
6. The degree of lysis will depend on tissue composition of the sample, as fibrotic or adipose tissue content can be variable. There may also be variation based on which muscle and what part of the muscle is biopsied. Additional lysis steps may be performed if large tissue fragments remain, however, the processing time prior to GEM generation should be minimized to avoid RNA degradation. Perform preliminary assessments to determine the number of lysis steps needed to isolate sufficient nuclei.

### Nuclear Isolation

7. Transfer remaining clear supernatant to the Lo-Bind tube
8. Centrifuge in a swinging bucket rotor at 500 g, 5 mins, 4°C. You should see a pellet containing mostly debris
9. Carefully pipet off the supernatant using a P200 to avoid disturbing the pellet
10. Add 200 µL Nuclei Wash and Resuspension buffer without resuspending, wait 5 minutes to allow buffer interchange
11. Using a cut 1000 µL tip, add 300 µL Nuclei Wash and Resuspension buffer and resuspend with 5 slow triturations
12. Filter the suspension through 40 µM FlowMi Cell Strainers into a labelled 5 mL round-bottom Falcon tube
13. Load 10 µL of suspension onto a hemocytometer, observe for nuclear quality and approximate number (Figure 1).

**Figure 1.**
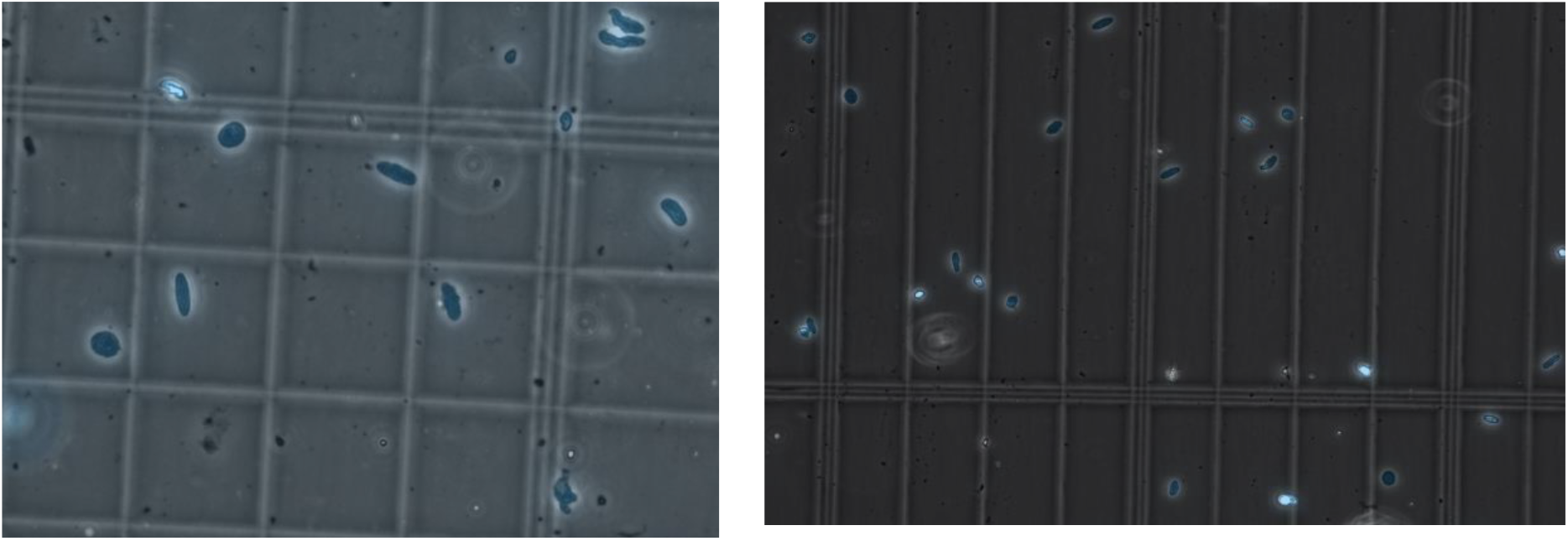
Representative images of nuclei quality before (A) and after (B) filtration and FANS sorting of nuclei. Nuclei are marked in blue by DAPI staining. Three images were taken for every sample, and each was observed at 40X magnification as recommended by 10X to look for nuclei integrity and the absence of clumping. After sorting, debris is reduced, nuclei are intact, and display minimal to no clumping.

### FANS Sorting

14. Sort the nuclei based on DAPI fluorescence into a tube with 100 µL of Nuc W+R and 4 U/µL RNase Inhibitor. This step is necessary because FANS sorting will dilute the Nuclei Wash and Resuspension Buffer, resulting in a low RNase Inhibitor concentration. Cells will be sitting on ice for extended periods of time without proper protection from RNA degradation ***RNase Inhibitor concentration assumes 2 mL final volume of sorted nuclei. Perform preliminary assessments to determine a suitable RNase Inhibitor concentration for sorting.
15. We gated nuclei by SSC-A/FSC-A for granularity and size. Singlets were then gated based on size P2 (FSC-H/FSC-A) and granularity P3 (SSC-H/SSC-A). Finally, the P4 population was sorted for Pacific-Blue positive cells
16. Transfer sorted nuclei to a Lo-Bind tube and centrifuge in a swinging bucket rotor at 500 g, 5 mins, 4°C
17. Carefully remove supernatant with a P200, leaving a small amount of buffer near the bottom to avoid losing nuclei
18. Resuspend slowly in 50-100 µL Nuclei Wash and Resuspension Buffer
19. Load 10 µL of suspension onto a hemocytometer, observe for nuclear quality and count to determine nuclei concentration

NOTE: At this stage, there should be minimal debris in solution with the nuclei. Ideally, nuclei should have an intact membrane and no blebbing. Ideal concentration ranges from 700-1200 nuclei/µL. If nuclei are too concentrated, dilute in more Nuc W+R buffer. If nuclei are not concentrated enough, centrifuge at 500 xg, 5 mins, 4°C and resuspend in a smaller volume. This will result in some nuclei loss. Alternatively, load a larger volume of nuclei suspension onto the 10X chromium device. The nuclei are now ready to proceed with 10X Genomics library preparation.

### Library Preparation and Sequencing

We adjusted the volume of nuclei suspension added to the 10X Genomics Chromium Controller to target 10,000 recovered nuclei. This number accounted for the expected ∼65% loss of nuclei during library preparation. Library preparation is detailed by 10X Genomics in Chromium Single Cell 3’ Reagent Kits User Guide (v3.1 Chemistry; Dual Index). Briefly, nuclei were suspended in gel emulsions (GEMs), barcoded, subjected to reverse transcription, and excess reagents are cleaned up. cDNA is amplified (12 cycles), quantified by TapeStation and assembled into a Chromium Single Cell 3′ Gene Expression Dual Index Library (14 amplification cycles), and SPRIselect beads used to size select the final cDNA libraries. Libraries were sequenced on the NovaSeq using SP 100 reagents to generate 800 million total read pairs. Read lengths were indicated by 10X Genomics as follows: read 1: 28, i7 index: 10 bp, i5: index 10 bp, and read 2: 90 bp.

### Library Analysis and Quality Control

Samples were demultiplexed using Cell Ranger v5.0.0. After demultiplexing, samples were aligned to a reference genome using Cell Ranger count, STAR alignment, and the reference transcriptome GRCh38-3.0.0. Because nuclei were the source of RNA, there is a high prevalence of pre-mRNA in the samples. Therefore, the “include-introns” option was used during Cell Ranger count.

Data processing, quality control, and analysis were performed in R using Seurat. Cells were filtered based on number of unique genes and mean reads per cell. Genes expressed in fewer than 3 cells were not included in the down-stream analyses. Cells with fewer than 50 genes or higher than 2% mitochondrial DNA content were also excluded, as they are likely dying or the membrane was ruptured. Outlier cells were excluded based on examination of n_FeatureRNA vs. n_CountRNA plots for combined samples. These measures serve to exclude doublets, artifacts, and poor-quality nuclei.

Samples were integrated following the Seurat vignette. Gene expression was log normalized and scaled. Dimensionality reduction was achieved by using the principal component that represented over 90% of variation, and less than 0.1% variation from the previous principal component. Nuclei populations and subpopulations were identified using canonical marker genes identified in the literature.

## 3. Results

### Consistent Quality Control on a Variety of Samples

Many sequencing, mapping, and cell metrics were consistently high across all samples (**Table 3**). All sequencing metrics exhibited little variation between samples. The average fraction of reads in cells was high (83%), suggesting low ambient RNA in the sample. This agrees with our microscopy observations, which demonstrated intact nuclei with little to no blebbing and minimal clumping (Figure 1). All samples except for M1 exceed 10X Genomics recommended minimum 70% fraction reads in cell. All samples meet 10X Genomics recommended minimum 20,000 mean reads per cell and recommended greater than 30% reads mapped confidently to the transcriptome. We observed a large number of reads corresponding to intronic regions (54%), however, this was expected because we are sequencing nuclear RNA. Although we targeted 10,000 nuclei, the number of recovered nuclei after sequencing varied. Our two controls recovered around 10,000 nuclei, with the disease samples yielding less. Although quality control metrics are rarely reported in full, our values were similar to previous studies (Barruet et al., 2020; Dos Santos et al., 2020; Kim et al., 2020; Petrany et al., 2020). In summary, quality control metrics meet 10X Genomics’ recommendations and are in line with values from similar studies.

**Table 3.** Quality Control Metrics: Quality control data represents six samples across two separate experiments. The average and standard deviation for all metrics is represented.

| Sequencing Metrics |  |  |  |  |  |  |  |
| --- | --- | --- | --- | --- | --- | --- | --- |
| Sample | No. of Reads | Valid Barcodes (%) | Valid UMIs (%) | Sequencing saturation (%) | Q30 bases in Barcode (%) | Q30 bases in RNA read (%) | Q30 Bases in UMI (%) |
| C1 | 229,291,982 | 96.7 | 99.9 | 72.7 | 96.4 | 93.7 | 96.2 |
| C2 | 268,498,276 | 96.2 | 100 | 36.9 | 95.2 | 92.1 | 94.9 |
| m1 | 243,061,000 | 96.6 | 99.9 | 79 | 96.3 | 93.1 | 96.2 |
| M1 | 110,311,933 | 95.2 | 99.9 | 91.8 | 96.4 | 94.5 | 96.3 |
| M2 | 252,256,865 | 97.3 | 99.9 | 88.6 | 96.5 | 94 | 96.4 |
| M3 | 226,248,973 | 96.2 | 100 | 37.7 | 95 | 91.9 | 94.7 |
| MEAN | 221,611,504.8 | 96.4 | 99.9 | 67.8 | 96.0 | 93.2 | 95.8 |
| STDV | 56,684,528.4 | 0.7 | 0.1 | 24.6 | 0.7 | 1.0 | 0.8 |

| Read Mapping Metrics |  |  |  |  |  |  |  |
| --- | --- | --- | --- | --- | --- | --- | --- |
| Sample | Mapped to genome (%) | mapped confidently to genome (%) | mapped confidently to intergenic regions (%) | mapped confidently to intronic regions (%) | mapped confidently to exonic regions (%) | mapped confidently to transcriptome (%) | mapped antisense to gene (%) |
| C1 | 96.8 | 92.9 | 6.2 | 54.8 | 31.9 | 69.4 | 15.9 |
| C2 | 92.5 | 87.7 | 7 | 61.3 | 19.4 | 44.2 | 35.4 |
| m1 | 96.7 | 93.3 | 6.3 | 56.3 | 30.6 | 67.8 | 17.7 |
| M1 | 92.9 | 65.1 | 6.6 | 40.1 | 18.3 | 52.8 | 4.1 |
| M2 | 96.2 | 91.4 | 7.5 | 54.8 | 29.1 | 72.6 | 9.4 |
| M3 | 91.4 | 86 | 7.2 | 58.9 | 19.9 | 44 | 33.7 |
| MEAN | 94.4 | 86.1 | 6.8 | 54.4 | 24.9 | 58.5 | 19.4 |
| STDV | 2.4 | 10.7 | 0.5 | 7.4 | 6.3 | 13.0 | 12.7 |

| Cell Metrics |  |  |  |  |  |  |
| --- | --- | --- | --- | --- | --- | --- |
| Sample | Recovered Nuclei | Fractions reads in cell (%) | Mean reads per cell | Median genes per cell | Total genes detected | Median UMI counts per cell |
| C1 | 9,697 | 89.4 | 23,345 | 1409 | 26,472 | 3232 |
| C2 | 13,030 | 90.8 | 20,565 | 1,981 | 28,120 | 4,729 |
| m1 | 8,531 | 91.1 | 28,279 | 1344 | 26,198 | 3008 |
| M1 | 1161 | 60.6 | 87,914 | 771 | 19,440 | 1221 |
| M2 | 3,944 | 79.1 | 61,406 | 1533 | 26,062 | 2730 |
| M3 | 6,997 | 86.9 | 32,183 | 2,469 | 28,497 | 6,654 |
| MEAN | 7,226.7 | 83.0 | 42,282.0 | 1,584.5 | 25,798.2 | 3,595.7 |
| STDV | 4,221.5 | 11.8 | 26,744.4 | 581.8 | 3,279.8 | 1,871.0 |

### Cell Populations Are Consistent with Previous Findings

In total, we recovered 43,325 nuclei from 6 skeletal muscle biopsies. We performed unbiased clustering on all nuclei (**Figure 2**). We observed read and mitochondrial DNA distribution throughout each cluster indicating that clustering was not strongly influenced by the number of reads per cell or mitochondrial DNA content (Figure S2).

**Figure 2.**
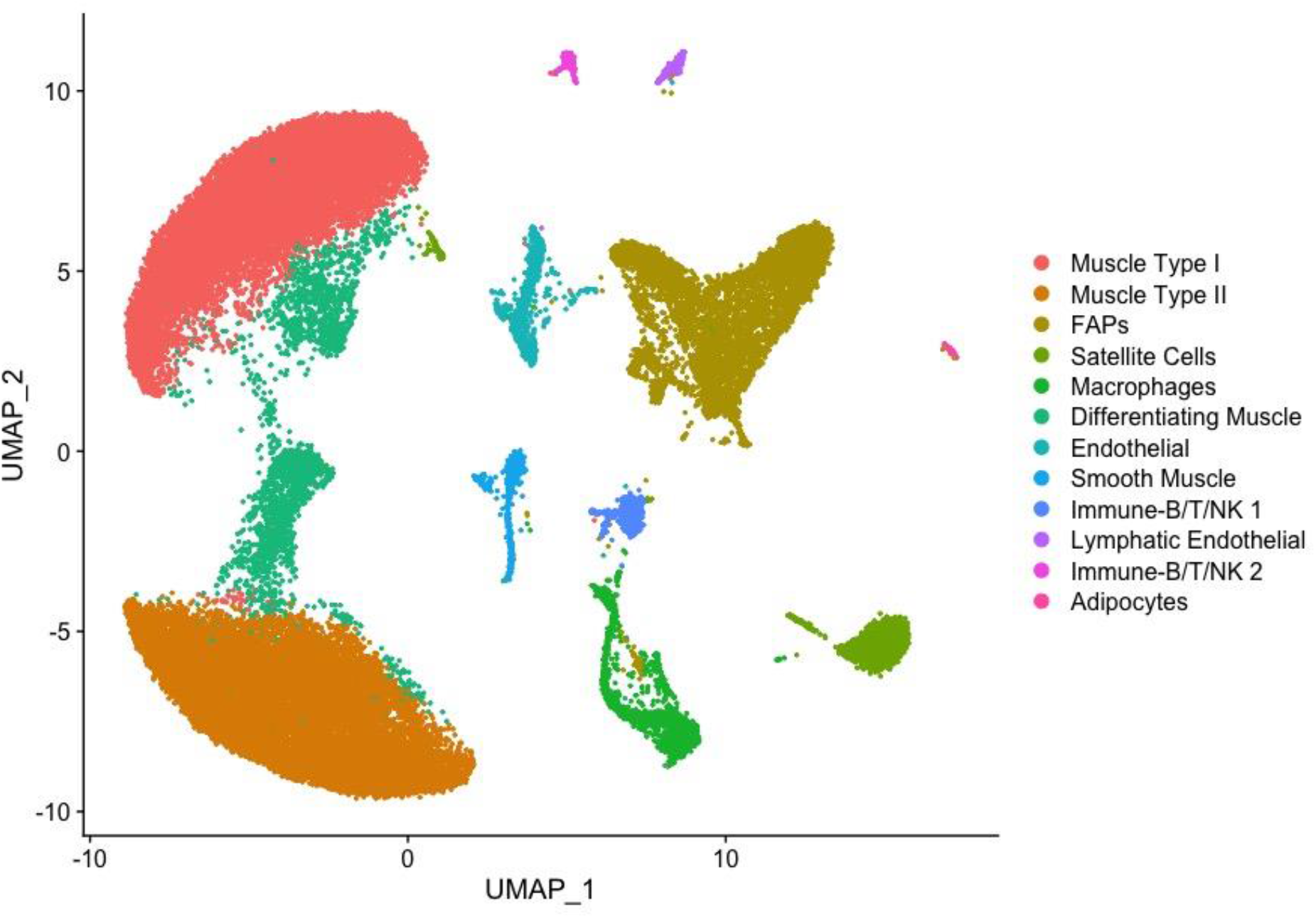
Marker Genes Used to Assign Cell Type. UMAP showing 43,325 nuclei separated by unbiased clustering.

We demonstrated that this method recovers cell populations expected in skeletal muscle (**Figure 2**). Canonical markers were used to assign the identity of each cluster (**Figure 3**). Overall, we identified 12 cell types whose identities are consistent with previous findings (De Micheli et al., 2020; Orchard et al., 2021; Rubenstein et al., 2020). For assignment of Type I and Type II fibers, we used previously reported gene sets (Rubenstein et al., 2020), which clearly separated Type I and Type II fibers (**Figure 4**). Genes reported by Rubenstein et al. to be differentially expressed between Type I and Type II fibers were highly correlated to our values for the same genes (Figure S5). We observed a third type of muscle in every sample which expressed both Type I and Type II muscle genes. Based on expression of genes only present in differentiating muscle like NCAM1 (Capkovic, Stevenson, Johnson, Thelen, & Cornelison, 2008), COL19A1 (Sumiyoshi, Laub, Yoshioka, & Ramirez, 2001), and MYH3 (Beylkin, Allen, & Leinwand, 2006), we concluded that these represent differentiating myonuclei.

**Figure 3.**
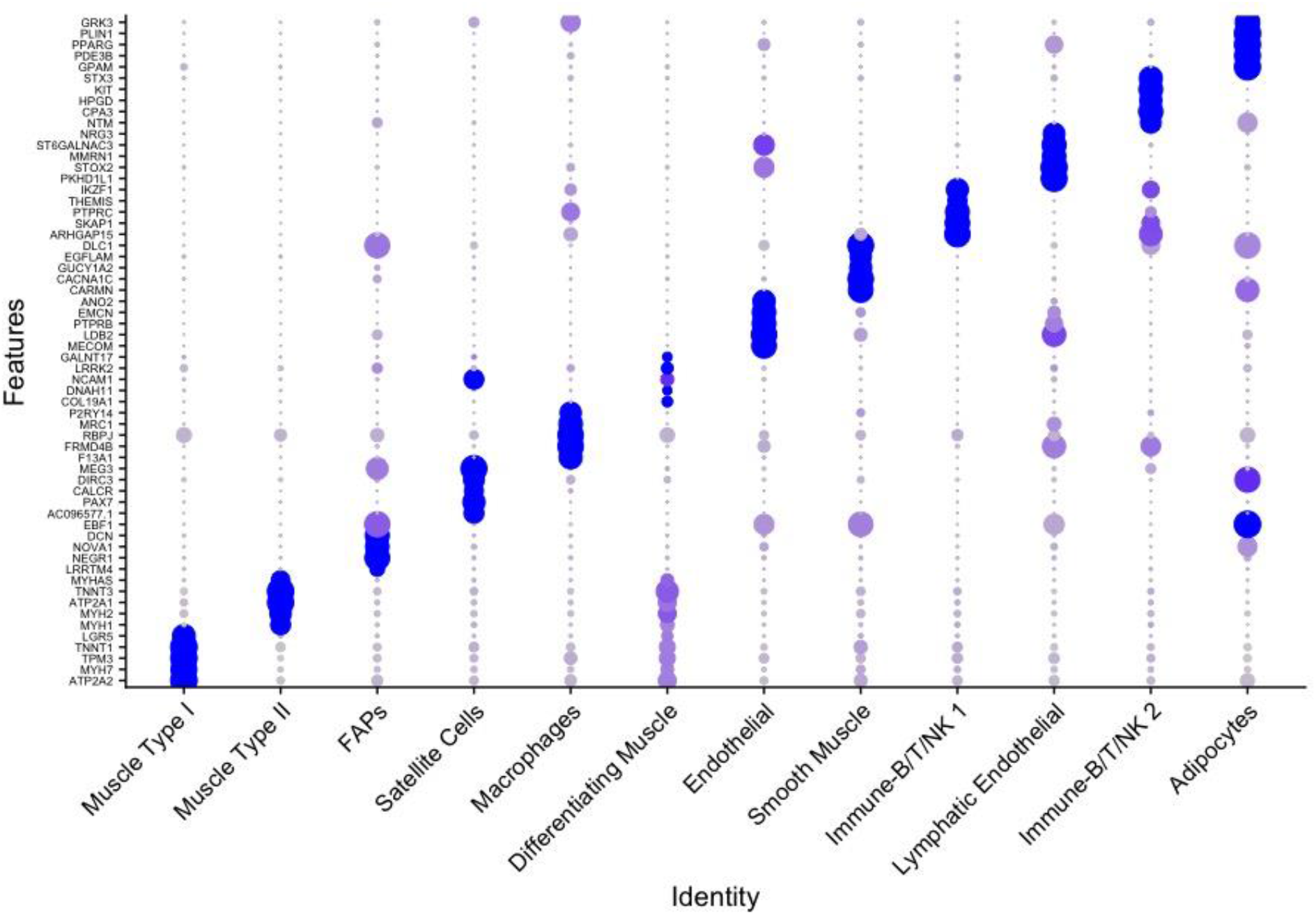
Marker Genes Used to Assign Cell Type. Dot Plot showing expression of top 5 markers for each cell type

**Figure 4.**
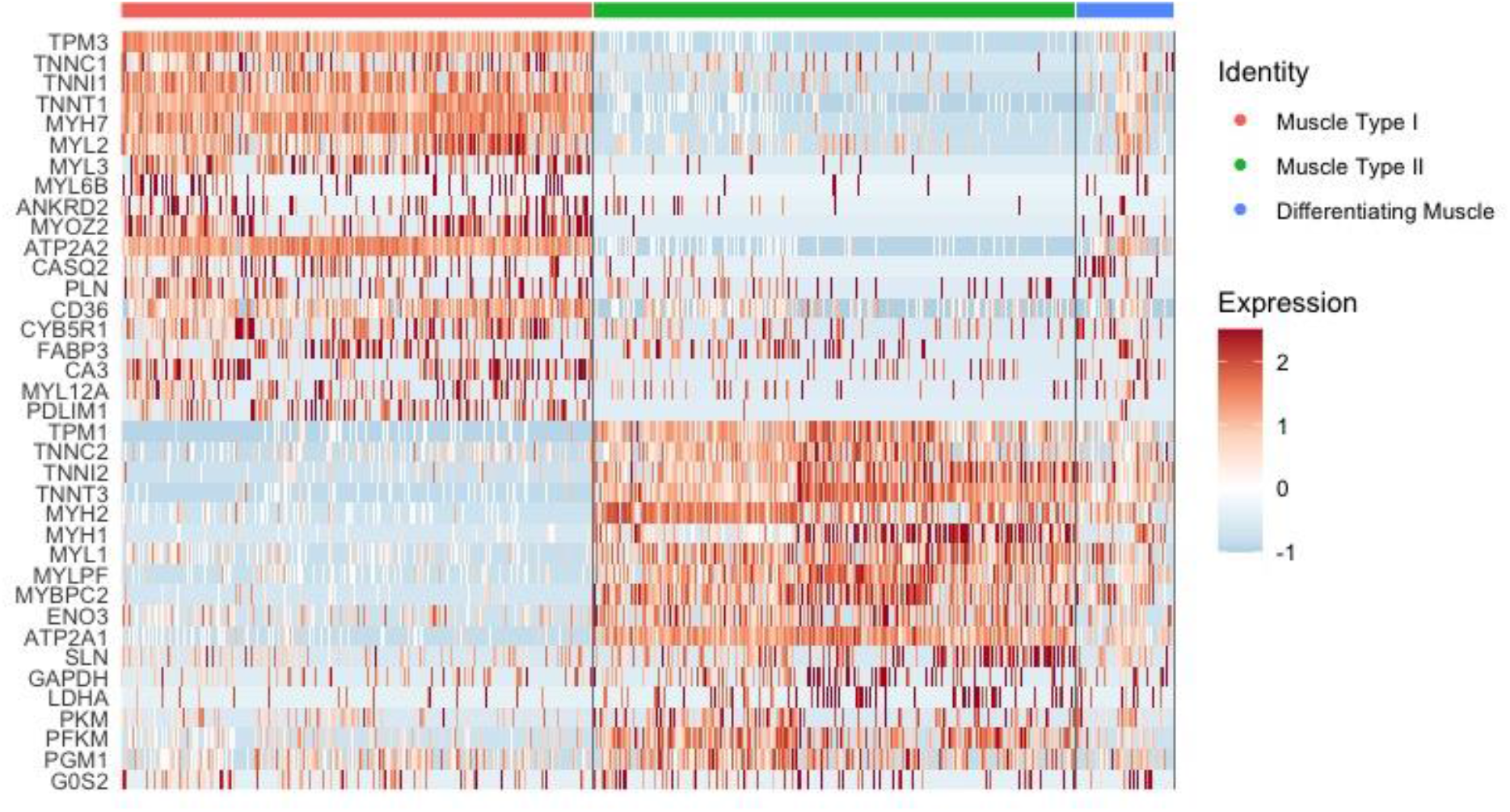
Marker Genes Used to Assign Cell Type. Heat map using genes identified by (Rubenstein et al., 2020) to distinguish type I and type II fibers.

Cell type proportions were very similar between controls C1 and C2, despite originating from different anatomical locations (Figure S3). Interestingly, samples m1 and M1 also closely matched the recovered cell type proportions of the controls. Cell type proportions in M2 and M3 varied from the controls and from each other. This was expected from samples with major histopathologic changes. Also, M2 and M3 still contained the same cell types, just in different proportions. On average, myonuclei and FAPs represented 82% of all nuclei. All other cell types made up a small proportion of the total cell number and displayed minor variations between samples.

A previous report found a population of lymphatic ECs common to multiple organs in mice, including muscle (Feng, Chen, Nguyen, Wu, & Li, 2019). Gene pathway analysis of the top 200 marker genes shows significant enrichment of lymphatic endothelial cell differentiation, blood vessel endothelial cell differentiation, and lymphangiogenesis (Figure S4). We conclude that this cluster represents lymphatic ECs.

## 4. Discussion

Here, we have demonstrated a practical and effective method of isolating nuclei from human skeletal muscle. Our method yields a sufficient quantity of nuclei for snRNAseq. In total, 43,325 nuclei were recovered. With this method, we identified 12 distinct cell types which are consistent with previous reports in human skeletal muscle (De Micheli et al., 2020; Orchard et al., 2021; Rubenstein et al., 2020). Notably, this included a large proportion of type I and type II myonuclei. To our knowledge, successful isolation of human myonuclei in significant quantities has only been reported once before (Orchard et al., 2021).

Importantly, we were able to obtain consistent quality control metrics from a variety of samples. This included muscle representing both sexes, collected from the commonly biopsied deltoid and vastus lateralis muscles, and from patients 33 up to 64 years of age. Additionally, the biopsy from patient M2 was performed in 2006 and still yielded 4000 nuclei (**Table 3**), suggesting that this method is robust to extended periods of freezing as is typical of banked samples. We included four distinct muscle disease samples to test the applicability of this method to studying muscle disease. Our results suggest that this method is a good choice for studying banked samples and rare muscle diseases, where maximizing information gained from the sample is key.

**Table 4** illustrates several advantages to our method. Specifically, ceramic beads avoid the difficulties associated with Dounce homogenization. Fluorescence assisted nuclei sorting (FANS) was used to purify nuclei, minimizing impact on RNA expression or integrity compared to enzymatic dissociation, which can affect transcription in single cells (Orchard et al., 2021; Van Den Brink et al., 2017). Overall, we have developed a method that minimizes the time between sample homogenization and gel in bead emulsion (GEM) generation, without sacrificing RNA integrity.

**Table 4.** Method Comparison. Justification of technical and practical advantages to our methodology.

| Previous Methods | Our Method | References |
| --- | --- | --- |
| **scRNAseq**<br>Requires fresh tissue<br>Large, multinucleated myofibers cannot pass through microfluidics<br>Very few myofibers represented | **snRNAseq**<br>Allows analysis of frozen banked samples<br>Nuclei from all cell types can pass through microfluidics<br>High proportion of myonuclei present | (De Micheli et al., 2020; Dos Santos et al., 2020; Slyper et al., 2020; Zeng et al., 2016) |
| **Enzymatic Dissociation**<br>Can influence transcriptional behaviour of muscle satellite cells<br>Typically requires an hour of enzymatic action | **FANS**<br>Increase the number of nuclei obtained<br>Minimal, if any impact on library quality | (Chongtham et al., 2021; Orchard et al., 2021; Van Den Brink et al., 2017) |
| **Dounce Homogenization**<br>Human muscle is too fibrous, yields too much debris, and results in poor filtering capacity and nuclei yields<br>Relatively slow process | **Lysing Using a Bead Tube**<br>Ceramic beads and 5-second pulses of high rpm shaking break down fibrous material<br>Relatively fast process | ‘Frankenstein’ protocol for nuclei isolation from fresh and frozen tissue |

Notably, we did not find a strong correlation between mass of homogenized muscle and quantity of recovered nuclei (Figure S1), suggesting either sample-specific effects or an effect of the ratio of lysis buffer to muscle mass. Regardless, we observed that masses of around 55-70 mg were ideal to avoid excess fibrous tissue that interfered with pipetting and filtering (Video 1). In addition to filtering, FANS was an effective method to get rid of excess debris that could interfere with downstream processing and microfluidics.

Two samples recovered significantly less nuclei than controls. We speculate that sample integrity could be compromised due to dead and dying cells already present at the time of tissue collection. This was illustrated by sample M2, which demonstrated a slightly lower percentage of reads in cells (79.1%), and less overall recovered nuclei (3,944) (**Table 3**). Sample M2 was biopsied in 2006, so freezing time could also play a role. Although quality control was similar to our other samples, further study is required to determine if age of muscle biopsy affects nuclei yield, or if insufficient nuclei were released during homogenization. Additionally, sample M1 had 60% of reads in cells and recovered 1,255 nuclei. A high reads per cell (87,914) and high sequencing saturation (91.8%) suggested a low complexity library. Notably, other QC metrics for both samples such as mean reads per cell, median genes per cell, and total genes detected were on par with controls. None of the samples underwent freeze thaw cycles before nuclei isolation, so this variable could not impact the quality of recovered nuclei. Additionally, it is unlikely that the isolation protocol induced a comparatively higher level of nuclei damage in these samples as they were processed side by side with other samples that recovered more nuclei. Overall, these results suggest that diseased samples with compromised tissue might reduce the total number of isolated nuclei, while still giving good quality data for the successfully recovered nuclei.

Although skeletal muscle contains type I, type IIa, type IId/x, and type IIb fibers (Hawley, Hargreaves, Joyner, & Zierath, 2014), we designated them either type I or type II. This is because we are aware of only one study that has characterized transcriptional differences between fiber types in human muscle (Rubenstein et al., 2020). Therefore, although we can only accurately distinguish between these two fiber types, it is obvious which nuclei represent type I and II muscle (**Figure 4**). Additionally, there was a high degree of correlation when comparing differentially expressed genes between Type I and Type II fibers identified by Rubenstein et al. (Figure S5). This gives confidence in accurate myonuclei assignments and demonstrates consistency between methods. Overall, a more detailed characterization of transcriptional variation between human muscle subtypes could lead to more accurate classification of snRNAseq derived cells and nuclei.

Proportions of Type I, Type II, and differentiating myofibers were relatively consistent between all samples except for M2 (Figure S3). Some variability between myofiber types was expected due to age (McCormick & Vasilaki, 2018), variable exercise regimes of participants (Hawley et al., 2014), and muscle specific differences. Additionally, selective atrophy of either fast or slow type muscle fibers has been reported for various muscle diseases (Wang & Pessin, 2013), as well as sarcopenia (Evans & Lexell, 1995). Other cell types also vary in number between people. For example, many factors influence the number of satellite cells in muscle tissue including age (W. Chen, Datzkiw, & Rudnicki, 2020; Yablonka-Reuveni, 2011), specific muscle location (Yablonka-Reuveni, 2011), fibre diameter (Maier & Bornemann, 1999), muscle type (I vs. IIa, IIx, and IIb) (Yin, Price, & Rudnicki, 2013), disease (Chang, Chevalier, & Rudnicki, 2016), inflammation, exercise and injury (W. Chen et al., 2020; Yablonka-Reuveni, 2011).

Additionally, the number of satellite cell nuclei recovered in the isolation procedure may depend on the extent of tissue mincing. Finally, variation between cell type proportions has been reported in single cell analysis of human muscles biopsied from different anatomical locations (De Micheli et al., 2020), and in mice at different ages (Petrany et al., 2020). Overall, care should be taken when interpreting cell type proportions present in a human muscle biopsy, as they are influenced by many factors.

Many cell types have been hypothesized to be involved in muscle disease, such as satellite cells in Duchenne muscular dystrophy (Chang et al., 2016), fiboradipogenic progenitors in limb girdle muscular dystrophy (Hogarth et al., 2019; Uezumi et al., 2011), and immune cells as integral parts of response to injury and muscle regeneration (B. Chen & Shan, 2019). However, we believe that examining myogenic cells is also crucial to gain a full understanding of the muscle environment. Importantly, we obtained many myonuclei using this technique, representing an average of 67% of our total nuclei. Interestingly, we were also able to identify two populations of ECs. Despite Feng and colleagues working with data from mouse ECs, 28% of their marker genes were also present in our lymphatic EC top marker genes, including MMRN1 and PROX1. Interestingly, like Feng and colleagues’ findings, MMRN1 was also a specific marker for lymphatic ECs in our human muscle samples (Figure S4). To our knowledge, this is the first report of lymphatic ECs in human muscle at the single cell level. These results emphasize this method’s ability to detect expected, as well as rare cell types.

Our method has the potential to elucidate transcriptional responses of muscle cell types in disease, exercise, and homeostatic conditions. Previously unreported in human single cell data, we identified the lymphatic EC. Although this emphasizes the robust nature of our approach, we only analyzed biopsies from two anatomical locations. Additional rare cell types could be uncovered in more diverse muscle biopsies. Future studies of banked samples with a variety of sexes, ages, and diseases would be valuable.

## Acknowledgements

TGBS is the recipient of a Hotchkiss Brain Institute Graduate Recruitment Scholarship. CSP is the recipient of an Eyes High Scholarship from the University of Calgary. SL is a recipient of Canada Graduate Scholarship from the Canadian Institutes of Health Research. Infrastructure for this work was supported by a John R Evans infrastructure grant from the Canada Foundation for Innovation to GP. MDN is supported by a CIHR project grant. Nuclei were sorted by the University of Calgary Flow Cytometry Core Facility. Libraries from nuclei were created by Dr. Rosin and the lab of Dr. Jeff Biernaskie. Libraries were sequenced at the University of Calgary Centre for Health Genomics and Informatics, which is supported by the Cumming School of Medicine. Cellranger was supported by resources from the Research Computing Services group at the University of Calgary.

## Ethics Statement

Work described in this study was approved by the University of Calgary Conjoint Health Research Ethics Board, REB16-2196.

## Others

https://www.protocols.io/view/frankenstein-protocol-for-nuclei-isolation-from-f-5jyl8nx98l2w/v3

