## Supplemental data for "A Protocol for Single Nucleus RNAseq from Frozen Skeletal Muscle"

**Supplementary Figures**

**Figure S1. No relationship between muscle mass and total nuclei recovered:** The number of nuclei recovered for analysis does not correlate with the muscle mass used for nuclei isolation. Each point represents one sample. R^2^ = 0.09

| Muscle Tissue (mg) | Recovered Nuclei |
| --- | --- |
| 72 | 7030 |
| 72 | 13056 |
| 94 | 1184 |
| 62 | 3949 |
| 65 | 8534 |
| 85 | 9700 |

**Figure S2. Cells cluster based on transcriptional profile:** Feature plots demonstrating that cells do not appear to cluster based on A) mitochondrial DNA or B) total transcripts per cell. Nuclei with more intense colour indicate increased percent mitochondrial DNA or transcripts per cell. There are no clusters or regions where nuclei exhibit a significantly higher number of transcripts. This indicates that nuclei are primarily clustering based on transcriptional profiles.

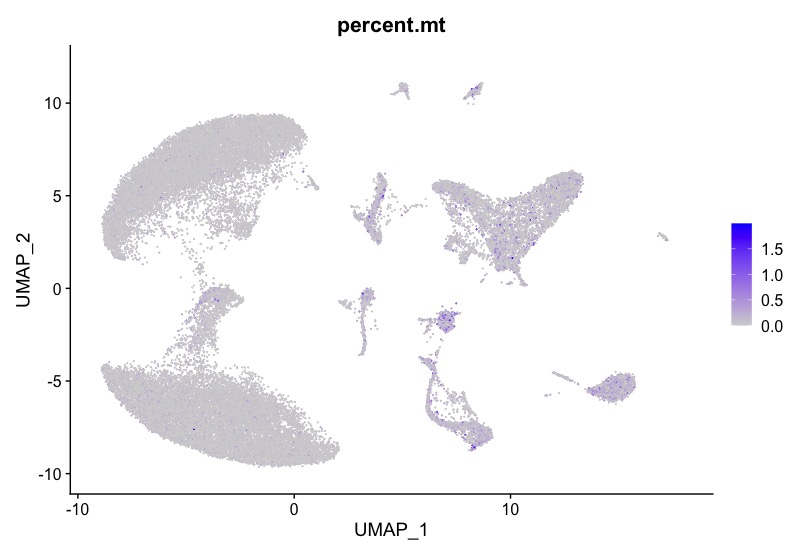

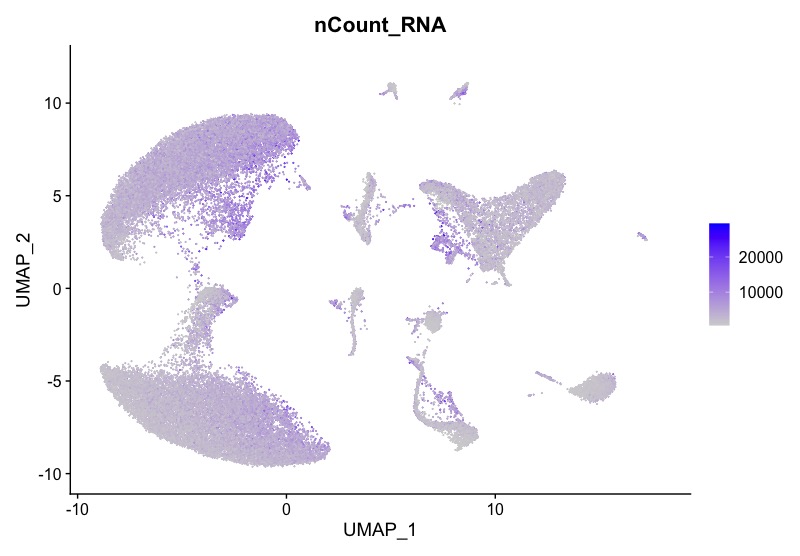

**Figure S3.** **Cell populations display consistent proportions:** Percentages of cells in each population. This represents three samples across two separate experiments.

Cell Type Proportion (%)

|  | Muscle Type I | Muscle Type II | FAPs | Macrophages | Satellite Cells | Differentiating Muscle | Endothelial | Smooth Muscle | B/T/NK 1 | Lymphatic Endothelial | B/T/NK 2 | Adipocytes |
| --- | --- | --- | --- | --- | --- | --- | --- | --- | --- | --- | --- | --- |
| C1 | 28.9398% | 41.3057% | 11.0045% | 4.67203% | 2.35149% | 6.51815% | 1.19637% | 2.05239% | 1.30982% | 0.0928218% | 0.556931% | 0% |
| C2 | 39.2053% | 34.0481% | 12.6355% | 2.42103% | 3.4125% | 3.82753% | 1.06064% | 2.39029% | 0.338175% | 0.276689% | 0.207517% | 0.176773% |
| m1 | 21.8146% | 46.6885% | 7.6896% | 7.99437% | 3.58692% | 5.98992% | 1.20736% | 0.773649% | 2.83671% | 0% | 1.41836% | 0% |
| M1 | 40.4145% | 31.0881% | 2.41796% | 3.97237% | 0.690846% | 18.3938% | 1.29534% | 0.777202% | 0.172712% | 0.518135% | 0% | 0.259067% |
| M2 | 5.70631% | 9.40908% | 37.4588% | 2.2318% | 10.804% | 11.1337% | 15.5719% | 1.16662% | 0.836926% | 3.82957% | 0% | 1.85138% |
| M3 | 43.6874% | 9.08961% | 16.1036% | 7.32894% | 8.71743% | 6.4844% | 1.2024% | 1.54595% | 2.27598% | 1.90381% | 0.672774% | 0.98769% |

**Figure S4. Lymphatic endothelial cells:** A) GO ontology of the top 200 marker genes for the lymphatic EC cluster. [Panther GO analysis](http://geneontology.org/) was used to generate these terms. B) MMRN1 is a specific marker of lymphatic endothelial cells.

-log10(P-value)

|  | *-LOG10(P-value) |
| --- | --- |
| lymphatic endothelial cell differentiation | 3.92812 |
| blood vessel endothelial cell differentiation | 3.6968 |
| endocardium development | 3.50031 |
| lymphangiogenesis | 3.41341 |
| neurotransmitter-gated ion channel clustering | 3.41341 |

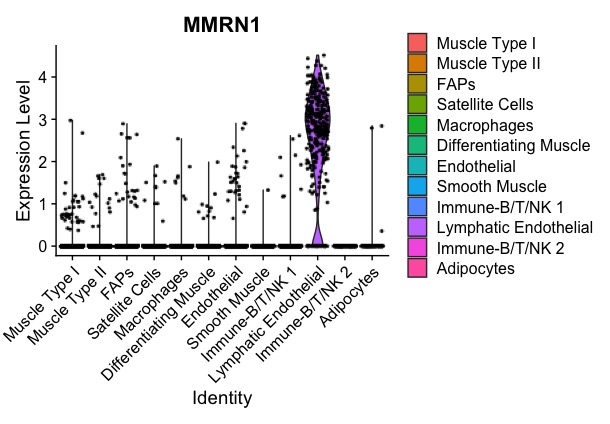

**Figure S5. Muscle marker genes are consistent with previous findings.** Log2(fold change) of type I and type II myofibers from Rubenstein et al. are compared with our values for the same genes. Of the top 40 differentially expressed genes found by Rubenstein et al., 35 were also differentially expressed between our type I and II myonuclei. The correlation between both sets of log2(FC) values is R^2^=0.85, however, for illustrative purposes we show a perfect line.

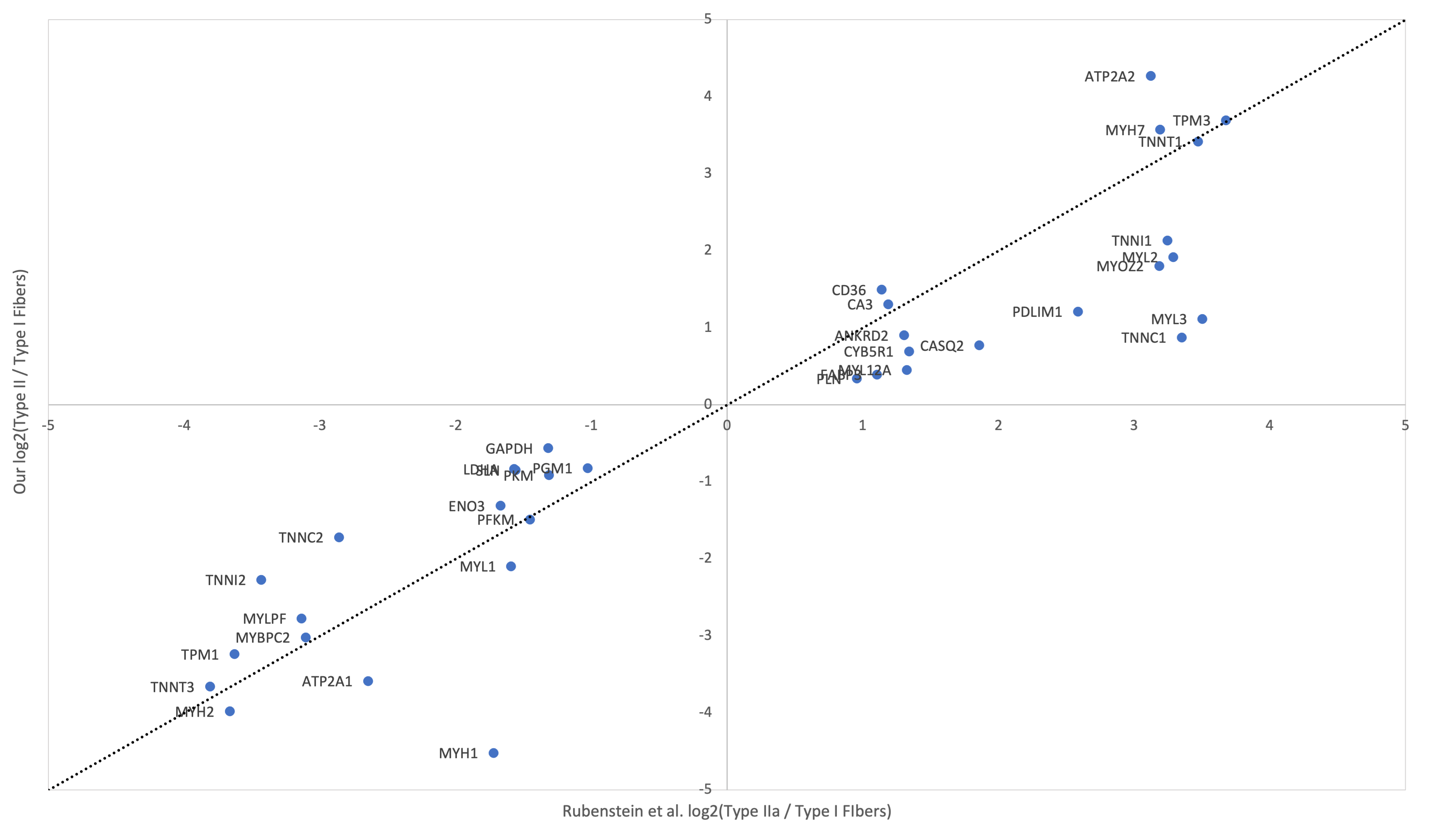

**Figure S6. Top 50 differentially expressed genes.** Differentially expressed genes were identified for each cluster using FindAllMarkers. The top 50 genes by log2(fold change) are reported, along with their fold change values, percentage of cells expressing the gene in Type I and Type II myonuclei, and adjusted p-value.

| avg_log2FC | pct.1 | pct.2 | p_val_adj | cluster | gene |
| --- | --- | --- | --- | --- | --- |
| 3.37309107 | 0.993 | 0.329 | 0 | Muscle Type I | ATP2A2 |
| 3.02079453 | 0.967 | 0.196 | 0 | Muscle Type I | MYH7 |
| 2.93937582 | 0.987 | 0.261 | 0 | Muscle Type I | TPM3 |
| 2.93320351 | 0.995 | 0.315 | 0 | Muscle Type I | TNNT1 |
| 2.57924759 | 0.843 | 0.13 | 0 | Muscle Type I | LGR5 |
| 2.29337341 | 0.823 | 0.076 | 0 | Muscle Type I | MYH7B |
| 2.26546446 | 0.949 | 0.386 | 0 | Muscle Type I | XPO4 |
| 1.99867785 | 0.681 | 0.058 | 0 | Muscle Type I | TECRL |
| 1.93336118 | 0.886 | 0.205 | 0 | Muscle Type I | TNNI1 |
| 1.74855472 | 0.84 | 0.256 | 0 | Muscle Type I | MYL2 |
| 1.74586286 | 0.802 | 0.213 | 0 | Muscle Type I | ESRRG |
| 1.73662404 | 0.847 | 0.205 | 0 | Muscle Type I | MYOM3 |
| 1.64348375 | 0.664 | 0.131 | 0 | Muscle Type I | ZNF385B |
| 1.60048147 | 0.49 | 0.062 | 0 | Muscle Type I | MYOZ2 |
| 1.40813378 | 0.914 | 0.611 | 0 | Muscle Type I | RIF1 |
| 1.38405268 | 0.906 | 0.396 | 0 | Muscle Type I | CD36 |
| 1.36142995 | 0.913 | 0.467 | 0 | Muscle Type I | USP54 |
| 1.27897541 | 0.867 | 0.421 | 0 | Muscle Type I | USP13 |
| 1.26564657 | 0.621 | 0.149 | 0 | Muscle Type I | ALPK2 |
| 1.26357013 | 0.411 | 0.073 | 0 | Muscle Type I | CCR3 |
| 1.25020171 | 0.504 | 0.056 | 0 | Muscle Type I | MYLK3 |
| 1.22704469 | 0.999 | 0.82 | 0 | Muscle Type I | NEB |
| 1.22515039 | 0.585 | 0.113 | 0 | Muscle Type I | SLC1A3 |
| 1.21849655 | 0.87 | 0.424 | 0 | Muscle Type I | RCAN2 |
| 1.21722634 | 0.459 | 0.134 | 0 | Muscle Type I | CA3 |
| 1.21569324 | 0.612 | 0.24 | 0 | Muscle Type I | GAS2 |
| 1.19240221 | 0.87 | 0.398 | 0 | Muscle Type I | TPD52L1 |
| 1.17701808 | 0.994 | 0.642 | 0 | Muscle Type I | MYOM1 |
| 1.17447382 | 0.298 | 0.028 | 0 | Muscle Type I | AC099066.2 |
| 1.17366624 | 0.516 | 0.123 | 0 | Muscle Type I | ASB18 |
| 1.14626553 | 0.749 | 0.384 | 0 | Muscle Type I | CRIM1 |
| 1.14374339 | 0.761 | 0.307 | 0 | Muscle Type I | LMOD2 |
| 1.14179642 | 0.841 | 0.448 | 0 | Muscle Type I | EXOC6 |
| 1.13445479 | 0.819 | 0.38 | 0 | Muscle Type I | AC106791.1 |
| 1.12348106 | 0.833 | 0.372 | 0 | Muscle Type I | KCNMA1 |
| 1.10462983 | 0.824 | 0.345 | 0 | Muscle Type I | LRRC39 |
| 1.1041716 | 0.582 | 0.21 | 0 | Muscle Type I | NEK10 |
| 1.09926639 | 0.555 | 0.125 | 0 | Muscle Type I | ATP1B4 |
| 1.09738811 | 0.904 | 0.436 | 0 | Muscle Type I | LRRC2 |
| 1.09291407 | 0.953 | 0.668 | 0 | Muscle Type I | TTN-AS1 |
| 1.08000308 | 0.988 | 0.602 | 0 | Muscle Type I | MYOT |
| 1.07407833 | 0.595 | 0.173 | 0 | Muscle Type I | PLIN5 |
| 1.07381726 | 0.838 | 0.366 | 0 | Muscle Type I | CKMT2 |
| 1.07273603 | 0.853 | 0.448 | 0 | Muscle Type I | SYNPO2 |
| 1.06564114 | 0.978 | 0.556 | 0 | Muscle Type I | ACTN2 |
| 1.05694305 | 0.515 | 0.072 | 0 | Muscle Type I | MYL3 |
| 1.05498063 | 0.999 | 0.769 | 0 | Muscle Type I | MYBPC1 |
| 1.04299413 | 0.983 | 0.716 | 0 | Muscle Type I | SGCD |
| 1.04055793 | 0.665 | 0.259 | 0 | Muscle Type I | P4HA1 |
| 1.03384749 | 0.299 | 0.052 | 0 | Muscle Type I | NKAIN3 |
| 3.60198318 | 0.731 | 0.135 | 0 | Muscle Type II | MYH1 |
| 3.21586573 | 0.792 | 0.251 | 0 | Muscle Type II | MYH2 |
| 3.17167175 | 0.976 | 0.225 | 0 | Muscle Type II | ATP2A1 |
| 2.87240991 | 0.986 | 0.271 | 0 | Muscle Type II | TNNT3 |
| 2.50938067 | 0.678 | 0.111 | 0 | Muscle Type II | MYHAS |
| 2.50912071 | 0.807 | 0.124 | 0 | Muscle Type II | MYBPC2 |
| 2.32199773 | 0.865 | 0.269 | 0 | Muscle Type II | MYLPF |
| 2.11669669 | 0.81 | 0.304 | 0 | Muscle Type II | MYL1 |
| 1.9888197 | 0.481 | 0.076 | 0 | Muscle Type II | ATRNL1 |
| 1.98266026 | 0.911 | 0.343 | 0 | Muscle Type II | TPM1 |
| 1.91954084 | 0.827 | 0.212 | 0 | Muscle Type II | TNNI2 |
| 1.90299083 | 0.312 | 0.03 | 0 | Muscle Type II | LINC02107 |
| 1.74123873 | 0.623 | 0.239 | 0 | Muscle Type II | MYLK4 |
| 1.70192377 | 0.387 | 0.045 | 0 | Muscle Type II | ACTN3 |
| 1.6404976 | 0.473 | 0.173 | 0 | Muscle Type II | GGT7 |
| 1.57733609 | 0.714 | 0.257 | 0 | Muscle Type II | PFKM |
| 1.56552765 | 0.351 | 0.054 | 0 | Muscle Type II | AC112206.2 |
| 1.51525403 | 0.895 | 0.555 | 0 | Muscle Type II | DENND2C |
| 1.50713964 | 0.87 | 0.497 | 0 | Muscle Type II | RHOBTB1 |
| 1.48628326 | 0.724 | 0.309 | 0 | Muscle Type II | PFKFB1 |
| 1.43498499 | 0.85 | 0.415 | 0 | Muscle Type II | COL4A3 |
| 1.4105992 | 0.681 | 0.27 | 0 | Muscle Type II | ENO3 |
| 1.40906649 | 0.869 | 0.508 | 0 | Muscle Type II | KCNQ5 |
| 1.40472367 | 0.738 | 0.376 | 0 | Muscle Type II | EGF |
| 1.38714312 | 0.541 | 0.224 | 0 | Muscle Type II | MLF1 |
| 1.37756079 | 0.731 | 0.426 | 0 | Muscle Type II | NEDD4 |
| 1.37335496 | 0.995 | 0.797 | 0 | Muscle Type II | FILIP1L |
| 1.36216203 | 0.798 | 0.331 | 0 | Muscle Type II | TNNC2 |
| 1.34668161 | 0.522 | 0.221 | 0 | Muscle Type II | UNC13C |
| 1.34359965 | 0.423 | 0.12 | 0 | Muscle Type II | SH3RF2 |
| 1.33739239 | 0.869 | 0.537 | 0 | Muscle Type II | DEPTOR |
| 1.33352381 | 0.81 | 0.446 | 0 | Muscle Type II | ARHGAP6 |
| 1.31958247 | 0.915 | 0.586 | 0 | Muscle Type II | AGL |
| 1.31586037 | 0.467 | 0.177 | 0 | Muscle Type II | LANCL1-AS1 |
| 1.30458321 | 0.857 | 0.483 | 0 | Muscle Type II | SLC7A2 |
| 1.30236125 | 0.766 | 0.389 | 0 | Muscle Type II | ART3 |
| 1.29063759 | 0.866 | 0.551 | 0 | Muscle Type II | PHTF2 |
| 1.24839231 | 0.985 | 0.674 | 0 | Muscle Type II | PDLIM3 |
| 1.24466598 | 0.958 | 0.697 | 0 | Muscle Type II | SESN1 |
| 1.23530984 | 0.47 | 0.141 | 0 | Muscle Type II | ATP2B2 |
| 1.22344382 | 0.8 | 0.455 | 0 | Muscle Type II | PPP1R3A |
| 1.22304237 | 0.418 | 0.119 | 0 | Muscle Type II | GADL1 |
| 1.2150497 | 0.68 | 0.366 | 0 | Muscle Type II | PGM1 |
| 1.19433888 | 0.731 | 0.431 | 0 | Muscle Type II | NOS1 |
| 1.19332863 | 0.8 | 0.446 | 0 | Muscle Type II | MYOZ1 |
| 1.19293919 | 0.695 | 0.369 | 0 | Muscle Type II | PYGM |
| 1.19129409 | 0.562 | 0.274 | 0 | Muscle Type II | PHKA1 |
| 1.1892849 | 0.502 | 0.197 | 0 | Muscle Type II | PSTPIP2 |
| 1.16639799 | 0.658 | 0.306 | 0 | Muscle Type II | COL4A4 |
| 1.15726439 | 0.832 | 0.491 | 0 | Muscle Type II | AMPD1 |
| 5.36688541 | 0.521 | 0.053 | 0 | FAPs | LRRTM4 |
| 4.83644062 | 0.921 | 0.065 | 0 | FAPs | NEGR1 |
| 4.34885191 | 0.872 | 0.056 | 0 | FAPs | NOVA1 |
| 3.86689131 | 0.89 | 0.056 | 0 | FAPs | DCN |
| 3.73521936 | 0.936 | 0.087 | 0 | FAPs | EBF1 |
| 3.58523186 | 0.785 | 0.081 | 0 | FAPs | FBN1 |
| 3.4850202 | 0.616 | 0.054 | 0 | FAPs | COL15A1 |
| 3.46181543 | 0.416 | 0.026 | 0 | FAPs | SCN7A |
| 3.44216934 | 0.79 | 0.044 | 0 | FAPs | TNXB |
| 3.42135385 | 0.506 | 0.033 | 0 | FAPs | ROBO2 |
| 3.37311396 | 0.774 | 0.03 | 0 | FAPs | COL6A3 |
| 3.35794196 | 0.825 | 0.07 | 0 | FAPs | ABCA8 |
| 3.34843863 | 0.8 | 0.06 | 0 | FAPs | DCLK1 |
| 3.29192007 | 0.947 | 0.501 | 0 | FAPs | LAMA2 |
| 3.15078771 | 0.289 | 0.015 | 0 | FAPs | PDZRN4 |
| 3.12898395 | 0.763 | 0.067 | 0 | FAPs | PID1 |
| 3.07833665 | 0.781 | 0.031 | 0 | FAPs | EBF2 |
| 3.02072866 | 0.924 | 0.08 | 0 | FAPs | DLC1 |
| 2.95992138 | 0.644 | 0.106 | 0 | FAPs | KAZN |
| 2.88142565 | 0.649 | 0.077 | 0 | FAPs | SMOC2 |
| 2.81845929 | 0.514 | 0.029 | 0 | FAPs | KCND2 |
| 2.80881542 | 0.527 | 0.018 | 0 | FAPs | BMPER |
| 2.77185066 | 0.72 | 0.032 | 0 | FAPs | COL1A2 |
| 2.73227234 | 0.774 | 0.165 | 0 | FAPs | ABCA9 |
| 2.70699639 | 0.833 | 0.09 | 0 | FAPs | SOX5 |
| 2.65956578 | 0.713 | 0.029 | 0 | FAPs | VIT |
| 2.63357625 | 0.731 | 0.405 | 0 | FAPs | ABCA10 |
| 2.5733929 | 0.413 | 0.014 | 0 | FAPs | ADH1B |
| 2.57212057 | 0.52 | 0.115 | 0 | FAPs | SDK1 |
| 2.52690428 | 0.865 | 0.249 | 0 | FAPs | EGFR |
| 2.5186937 | 0.587 | 0.02 | 0 | FAPs | SVEP1 |
| 2.51710094 | 0.585 | 0.035 | 0 | FAPs | ADAMTSL3 |
| 2.50378848 | 0.491 | 0.081 | 0 | FAPs | MME |
| 2.47671695 | 0.382 | 0.021 | 0 | FAPs | NCAM2 |
| 2.45990423 | 0.521 | 0.012 | 0 | FAPs | MFAP5 |
| 2.43312014 | 0.741 | 0.107 | 0 | FAPs | HSPG2 |
| 2.42445823 | 0.627 | 0.096 | 0 | FAPs | CAMK1D |
| 2.42362522 | 0.422 | 0.011 | 0 | FAPs | NOX4 |
| 2.41113172 | 0.783 | 0.109 | 0 | FAPs | FBXL7 |
| 2.38482307 | 0.763 | 0.204 | 0 | FAPs | ABCA6 |
| 2.37799344 | 0.533 | 0.015 | 0 | FAPs | ITGA11 |
| 2.37628388 | 0.612 | 0.045 | 0 | FAPs | COL3A1 |
| 2.37448776 | 0.465 | 0.028 | 0 | FAPs | CCDC80 |
| 2.36976996 | 0.706 | 0.14 | 0 | FAPs | GSN |
| 2.32844281 | 0.643 | 0.08 | 0 | FAPs | UST |
| 2.32571923 | 0.618 | 0.076 | 0 | FAPs | GRK5 |
| 2.30475157 | 0.499 | 0.013 | 0 | FAPs | COL12A1 |
| 2.2938846 | 0.548 | 0.057 | 0 | FAPs | COL4A2 |
| 2.29132694 | 0.691 | 0.155 | 0 | FAPs | ABI3BP |
| 2.2899734 | 0.744 | 0.172 | 0 | FAPs | RUNX1T1 |
| 4.75250915 | 0.732 | 0.007 | 0 | Satellite Cells | AC096577.1 |
| 4.06508908 | 0.822 | 0.022 | 0 | Satellite Cells | PAX7 |
| 3.88753278 | 0.691 | 0.01 | 0 | Satellite Cells | CALCR |
| 3.79689447 | 0.765 | 0.137 | 0 | Satellite Cells | DIRC3 |
| 3.7508206 | 0.969 | 0.148 | 0 | Satellite Cells | MEG3 |
| 3.53892157 | 0.659 | 0.024 | 0 | Satellite Cells | TRHDE |
| 3.44319208 | 0.658 | 0.045 | 0 | Satellite Cells | CDH4 |
| 3.40336322 | 0.991 | 0.581 | 0 | Satellite Cells | CADM2 |
| 3.35092475 | 0.758 | 0.076 | 0 | Satellite Cells | GPC6 |
| 3.35041416 | 0.835 | 0.161 | 0 | Satellite Cells | CLCN5 |
| 3.28729208 | 0.77 | 0.091 | 0 | Satellite Cells | 01-Mar |
| 3.27910181 | 0.905 | 0.127 | 0 | Satellite Cells | HMCN2 |
| 3.25761115 | 0.536 | 0.013 | 0 | Satellite Cells | AC004053.1 |
| 3.24755219 | 0.802 | 0.099 | 0 | Satellite Cells | MEG8 |
| 3.1335308 | 0.709 | 0.043 | 0 | Satellite Cells | TENM4 |
| 2.70266598 | 0.571 | 0.04 | 0 | Satellite Cells | CNKSR3 |
| 2.69391644 | 0.742 | 0.188 | 0 | Satellite Cells | FRMD4A |
| 2.67354165 | 0.485 | 0.018 | 0 | Satellite Cells | TMEFF2 |
| 2.64252394 | 0.608 | 0.158 | 0 | Satellite Cells | KCNQ1OT1 |
| 2.63473516 | 0.727 | 0.09 | 0 | Satellite Cells | NCAM1 |
| 2.60430584 | 0.769 | 0.152 | 0 | Satellite Cells | DOCK9 |
| 2.5961996 | 0.428 | 0.005 | 0 | Satellite Cells | NLGN4X |
| 2.58635704 | 0.593 | 0.048 | 0 | Satellite Cells | DANT2 |
| 2.58233769 | 0.768 | 0.295 | 0 | Satellite Cells | KANK1 |
| 2.50430604 | 0.6 | 0.072 | 0 | Satellite Cells | MSC-AS1 |
| 2.47745033 | 0.79 | 0.318 | 0 | Satellite Cells | SPATS2L |
| 2.45427565 | 0.87 | 0.418 | 0 | Satellite Cells | TLN2 |
| 2.44271633 | 0.718 | 0.228 | 0 | Satellite Cells | PTCHD1-AS |
| 2.43261188 | 0.458 | 0.028 | 0 | Satellite Cells | GNA14 |
| 2.41831303 | 0.559 | 0.081 | 0 | Satellite Cells | NTN4 |
| 2.40407071 | 0.59 | 0.152 | 0 | Satellite Cells | DYNC1I1 |
| 2.33071535 | 0.604 | 0.129 | 0 | Satellite Cells | MEGF10 |
| 2.3046954 | 0.569 | 0.073 | 0 | Satellite Cells | PON2 |
| 2.30284589 | 0.509 | 0.033 | 0 | Satellite Cells | CHN1 |
| 2.26311237 | 0.51 | 0.063 | 0 | Satellite Cells | ITGBL1 |
| 2.24443552 | 0.703 | 0.247 | 0 | Satellite Cells | ADAMTS9-AS2 |
| 2.20122145 | 0.569 | 0.125 | 0 | Satellite Cells | PRKD1 |
| 2.19824795 | 0.64 | 0.124 | 0 | Satellite Cells | SPARCL1 |
| 2.19242704 | 0.518 | 0.086 | 0 | Satellite Cells | FN1 |
| 2.15291889 | 0.448 | 0.022 | 0 | Satellite Cells | OLFML2B |
| 2.14288412 | 0.88 | 0.339 | 0 | Satellite Cells | PLXDC2 |
| 2.109816 | 0.564 | 0.106 | 0 | Satellite Cells | RASSF4 |
| 2.0858242 | 0.487 | 0.041 | 0 | Satellite Cells | PXDN |
| 2.08342895 | 0.812 | 0.457 | 0 | Satellite Cells | PPP1R9A |
| 2.06002375 | 0.398 | 0.061 | 0 | Satellite Cells | LINC01239 |
| 2.02717126 | 0.373 | 0.007 | 0 | Satellite Cells | GRIK4 |
| 2.02122612 | 0.316 | 0.007 | 0 | Satellite Cells | CTNND2 |
| 2.01843641 | 0.619 | 0.135 | 0 | Satellite Cells | NAV1 |
| 2.01474014 | 0.426 | 0.045 | 0 | Satellite Cells | MUSK |
| 1.98450862 | 0.56 | 0.15 | 0 | Satellite Cells | MDFIC |
| 4.8822424 | 0.853 | 0.016 | 0 | Macrophages | F13A1 |
| 4.86871583 | 0.952 | 0.064 | 0 | Macrophages | FRMD4B |
| 4.46071449 | 0.947 | 0.452 | 0 | Macrophages | RBPJ |
| 4.26586887 | 0.872 | 0.017 | 0 | Macrophages | MRC1 |
| 3.83040802 | 0.787 | 0.019 | 0 | Macrophages | P2RY14 |
| 3.75775064 | 0.717 | 0.067 | 0 | Macrophages | COLEC12 |
| 3.68431779 | 0.797 | 0.069 | 0 | Macrophages | RGL1 |
| 3.62584249 | 0.851 | 0.026 | 0 | Macrophages | IQGAP2 |
| 3.62071212 | 0.795 | 0.146 | 0 | Macrophages | MAMDC2 |
| 3.60471483 | 0.821 | 0.014 | 0 | Macrophages | RBM47 |
| 3.55490999 | 0.789 | 0.197 | 0 | Macrophages | NAV2 |
| 3.47875888 | 0.678 | 0.07 | 0 | Macrophages | LGMN |
| 3.38074967 | 0.917 | 0.217 | 0 | Macrophages | SLC9A9 |
| 3.36657104 | 0.774 | 0.094 | 0 | Macrophages | MAN1A1 |
| 3.34909137 | 0.776 | 0.007 | 0 | Macrophages | MS4A6A |
| 3.33987856 | 0.751 | 0.029 | 0 | Macrophages | SCN9A |
| 3.26189938 | 0.749 | 0.061 | 0 | Macrophages | SLC8A1 |
| 3.25047661 | 0.722 | 0.009 | 0 | Macrophages | MS4A4E |
| 3.21594475 | 0.791 | 0.014 | 0 | Macrophages | TBXAS1 |
| 3.19281696 | 0.708 | 0.039 | 0 | Macrophages | DAB2 |
| 3.17841979 | 0.689 | 0.04 | 0 | Macrophages | HRH1 |
| 3.16629388 | 0.934 | 0.167 | 0 | Macrophages | LRMDA |
| 3.11522617 | 0.65 | 0.015 | 0 | Macrophages | LYVE1 |
| 3.06096613 | 0.79 | 0.027 | 0 | Macrophages | DOCK2 |
| 2.97431022 | 0.737 | 0.007 | 0 | Macrophages | SYK |
| 2.95085084 | 0.7 | 0.113 | 0 | Macrophages | SELENOP |
| 2.94480637 | 0.648 | 0.01 | 0 | Macrophages | STAB1 |
| 2.90940678 | 0.718 | 0.016 | 0 | Macrophages | ATP8B4 |
| 2.89124177 | 0.582 | 0.023 | 0 | Macrophages | RTN1 |
| 2.85547687 | 0.758 | 0.202 | 0 | Macrophages | PDGFC |
| 2.83220383 | 0.779 | 0.189 | 0 | Macrophages | NRP1 |
| 2.82996286 | 0.465 | 0.023 | 0 | Macrophages | LSAMP |
| 2.82590147 | 0.893 | 0.316 | 0 | Macrophages | ZEB2 |
| 2.73740569 | 0.662 | 0.026 | 0 | Macrophages | ADAP2 |
| 2.73509534 | 0.576 | 0.006 | 0 | Macrophages | CD163L1 |
| 2.70672425 | 0.751 | 0.266 | 0 | Macrophages | ITSN1 |
| 2.70374774 | 0.611 | 0.004 | 0 | Macrophages | LILRB5 |
| 2.673384 | 0.67 | 0.01 | 0 | Macrophages | CSF1R |
| 2.66126764 | 0.573 | 0.006 | 0 | Macrophages | SIGLEC1 |
| 2.63650113 | 0.58 | 0.026 | 0 | Macrophages | CPM |
| 2.62361991 | 0.647 | 0.014 | 0 | Macrophages | EMB |
| 2.60499806 | 0.72 | 0.1 | 0 | Macrophages | DAPK1 |
| 2.6040385 | 0.628 | 0.028 | 0 | Macrophages | SLCO2B1 |
| 2.59535616 | 0.778 | 0.287 | 0 | Macrophages | STARD13 |
| 2.58017583 | 0.627 | 0.093 | 0 | Macrophages | DNM1 |
| 2.57902682 | 0.605 | 0.004 | 0 | Macrophages | CD163 |
| 2.57280267 | 0.659 | 0.105 | 0 | Macrophages | AFF3 |
| 2.55051511 | 0.83 | 0.251 | 0 | Macrophages | ZSWIM6 |
| 2.52922507 | 0.686 | 0.104 | 0 | Macrophages | TNFAIP2 |
| 2.52819495 | 0.759 | 0.059 | 0 | Macrophages | FLI1 |
| 2.70701584 | 0.366 | 0.016 | 0 | Differentiating Muscle | COL19A1 |
| 2.13840535 | 0.298 | 0.033 | 0 | Differentiating Muscle | DNAH11 |
| 2.11234746 | 0.457 | 0.098 | 0 | Differentiating Muscle | NCAM1 |
| 2.08463021 | 0.408 | 0.198 | 1.53E-189 | Differentiating Muscle | LRRK2 |
| 1.84459552 | 0.316 | 0.048 | 0 | Differentiating Muscle | GALNT17 |
| 1.69807374 | 0.511 | 0.2 | 0 | Differentiating Muscle | ARHGAP28 |
| 1.63788677 | 0.503 | 0.204 | 0 | Differentiating Muscle | MDM2 |
| 1.63161351 | 0.261 | 0.08 | 8.34E-241 | Differentiating Muscle | COL21A1 |
| 1.49338812 | 0.338 | 0.099 | 0 | Differentiating Muscle | EFCAB7 |
| 1.48263946 | 0.386 | 0.174 | 1.03E-186 | Differentiating Muscle | RUNX1 |
| 1.46524838 | 0.515 | 0.241 | 2.14E-273 | Differentiating Muscle | ASTN2 |
| 1.4417495 | 0.416 | 0.149 | 0 | Differentiating Muscle | CASQ2 |
| 1.42796616 | 0.411 | 0.204 | 8.28E-158 | Differentiating Muscle | RNLS |
| 1.42142726 | 0.752 | 0.45 | 9.78E-287 | Differentiating Muscle | ARPP21 |
| 1.40425948 | 0.406 | 0.164 | 5.80E-265 | Differentiating Muscle | FAM184B |
| 1.36498797 | 0.755 | 0.49 | 1.44E-250 | Differentiating Muscle | OSBPL6 |
| 1.31904789 | 0.606 | 0.329 | 6.14E-261 | Differentiating Muscle | SLC7A6 |
| 1.30749605 | 0.332 | 0.139 | 3.96E-176 | Differentiating Muscle | AF165147.1 |
| 1.26888876 | 0.321 | 0.096 | 0 | Differentiating Muscle | PPP1R14C |
| 1.26731771 | 0.392 | 0.137 | 0 | Differentiating Muscle | ADAM23 |
| 1.26347321 | 0.358 | 0.154 | 3.59E-194 | Differentiating Muscle | FAM13C |
| 1.24436218 | 0.31 | 0.107 | 7.88E-237 | Differentiating Muscle | PLCE1 |
| 1.21309058 | 0.265 | 0.183 | 1.26E-23 | Differentiating Muscle | AL390957.1 |
| 1.20652115 | 0.96 | 0.734 | 0 | Differentiating Muscle | RBFOX1 |
| 1.19638069 | 0.682 | 0.383 | 1.56E-270 | Differentiating Muscle | BEST3 |
| 1.16740399 | 0.407 | 0.149 | 3.05E-285 | Differentiating Muscle | KCNN3 |
| 1.14903994 | 0.904 | 0.606 | 0 | Differentiating Muscle | TP63 |
| 1.13325873 | 0.802 | 0.622 | 3.90E-131 | Differentiating Muscle | XIRP2 |
| 1.12156099 | 0.405 | 0.151 | 2.00E-278 | Differentiating Muscle | FREM2 |
| 1.12118145 | 0.252 | 0.108 | 2.40E-121 | Differentiating Muscle | PRUNE2 |
| 1.10997857 | 0.618 | 0.425 | 1.80E-120 | Differentiating Muscle | GSE1 |
| 1.09326733 | 0.533 | 0.264 | 2.22E-231 | Differentiating Muscle | EPHB1 |
| 1.08022853 | 0.365 | 0.142 | 1.10E-232 | Differentiating Muscle | CCDC39.1 |
| 1.07553143 | 0.411 | 0.244 | 2.60E-98 | Differentiating Muscle | SH3PXD2A |
| 1.07201566 | 0.623 | 0.366 | 2.90E-187 | Differentiating Muscle | FLRT2 |
| 1.06395708 | 0.365 | 0.147 | 1.44E-208 | Differentiating Muscle | TMEM178B |
| 1.0631268 | 0.441 | 0.204 | 2.46E-211 | Differentiating Muscle | AMOTL1 |
| 1.03866605 | 0.616 | 0.358 | 1.83E-183 | Differentiating Muscle | SLC24A3 |
| 1.0275498 | 0.609 | 0.364 | 5.57E-172 | Differentiating Muscle | INPP4B |
| 1.00174695 | 0.491 | 0.291 | 1.35E-123 | Differentiating Muscle | ADARB1 |
| 1.00016953 | 0.325 | 0.099 | 1.59E-296 | Differentiating Muscle | CDC42EP3 |
| 0.99604536 | 0.538 | 0.378 | 1.92E-65 | Differentiating Muscle | LINC01091 |
| 0.98841721 | 0.391 | 0.179 | 1.99E-178 | Differentiating Muscle | RRAD |
| 0.94960273 | 0.613 | 0.378 | 3.64E-154 | Differentiating Muscle | PLA2G4C |
| 0.94745797 | 0.542 | 0.293 | 4.86E-189 | Differentiating Muscle | TRIM55 |
| 0.93980259 | 0.745 | 0.6 | 1.33E-84 | Differentiating Muscle | FOXO1 |
| 0.93886382 | 0.255 | 0.101 | 1.24E-140 | Differentiating Muscle | COBLL1 |
| 0.92835866 | 0.289 | 0.117 | 1.89E-156 | Differentiating Muscle | ITGA9 |
| 0.91477933 | 0.31 | 0.105 | 2.17E-237 | Differentiating Muscle | UCK2 |
| 0.90518116 | 0.607 | 0.403 | 3.75E-120 | Differentiating Muscle | TULP4 |
| 4.87576309 | 0.918 | 0.012 | 0 | Endothelial | MECOM |
| 4.35006148 | 0.945 | 0.072 | 0 | Endothelial | LDB2 |
| 4.08820924 | 0.882 | 0.01 | 0 | Endothelial | PTPRB |
| 4.03070028 | 0.879 | 0.016 | 0 | Endothelial | EMCN |
| 4.02243057 | 0.828 | 0.006 | 0 | Endothelial | ANO2 |
| 3.79025025 | 0.734 | 0.015 | 0 | Endothelial | SNTG2 |
| 3.73721442 | 0.822 | 0.011 | 0 | Endothelial | VWF |
| 3.68151574 | 0.754 | 0.03 | 0 | Endothelial | ST6GALNAC3 |
| 3.57157342 | 0.734 | 0.012 | 0 | Endothelial | FLT1 |
| 3.51167498 | 0.821 | 0.01 | 0 | Endothelial | EGFL7 |
| 3.51098134 | 0.827 | 0.038 | 0 | Endothelial | PECAM1 |
| 3.49281801 | 0.651 | 0.007 | 0 | Endothelial | TPO |
| 3.43093673 | 0.747 | 0.008 | 0 | Endothelial | CYYR1 |
| 3.34910781 | 0.776 | 0.24 | 0 | Endothelial | ARL15 |
| 3.2537259 | 0.749 | 0.062 | 0 | Endothelial | MCTP1 |
| 3.13325884 | 0.457 | 0.003 | 0 | Endothelial | BTNL9 |
| 3.03468636 | 0.343 | 0.011 | 0 | Endothelial | FAM155A |
| 3.03384572 | 0.821 | 0.073 | 0 | Endothelial | FLI1 |
| 3.01888844 | 0.797 | 0.147 | 0 | Endothelial | PLCB4 |
| 3.00033135 | 0.754 | 0.013 | 0 | Endothelial | ERG |
| 2.98713072 | 0.722 | 0.109 | 0 | Endothelial | PKP4 |
| 2.98290598 | 0.775 | 0.105 | 0 | Endothelial | PITPNC1 |
| 2.9715622 | 0.702 | 0.058 | 0 | Endothelial | ITGA6 |
| 2.9267261 | 0.72 | 0.014 | 0 | Endothelial | SHANK3 |
| 2.92326169 | 0.639 | 0.003 | 0 | Endothelial | ADGRL4 |
| 2.89321145 | 0.564 | 0.04 | 0 | Endothelial | CCDC85A |
| 2.87173446 | 0.744 | 0.133 | 0 | Endothelial | TSHZ2 |
| 2.86491623 | 0.649 | 0.006 | 0 | Endothelial | CXorf36 |
| 2.85254857 | 0.53 | 0.016 | 0 | Endothelial | DACH1 |
| 2.84795505 | 0.729 | 0.078 | 0 | Endothelial | STOX2 |
| 2.8434149 | 0.612 | 0.016 | 0 | Endothelial | RASGRF2 |
| 2.83893952 | 0.756 | 0.049 | 0 | Endothelial | ARHGAP31 |
| 2.83608602 | 0.519 | 0.042 | 0 | Endothelial | THSD7A |
| 2.78948529 | 0.684 | 0.052 | 0 | Endothelial | EPAS1 |
| 2.75940312 | 0.76 | 0.09 | 0 | Endothelial | PREX2 |
| 2.75878536 | 0.651 | 0.033 | 0 | Endothelial | PLEKHG1 |
| 2.71208827 | 0.983 | 0.734 | 0 | Endothelial | PTPRM |
| 2.70980362 | 0.56 | 0.116 | 0 | Endothelial | MYRIP |
| 2.69967462 | 0.669 | 0.053 | 0 | Endothelial | CSGALNACT1 |
| 2.6932681 | 0.408 | 0.014 | 0 | Endothelial | TLL1 |
| 2.6802064 | 0.551 | 0.066 | 0 | Endothelial | ABLIM3 |
| 2.64032044 | 0.679 | 0.178 | 0 | Endothelial | ZNF385D |
| 2.63725015 | 0.928 | 0.184 | 0 | Endothelial | LRMDA |
| 2.62710733 | 0.484 | 0.018 | 0 | Endothelial | ADGRF5 |
| 2.61122055 | 0.594 | 0.095 | 0 | Endothelial | MGLL |
| 2.6065919 | 0.892 | 0.173 | 0 | Endothelial | ELMO1 |
| 2.54376369 | 0.607 | 0.054 | 0 | Endothelial | GALNT18 |
| 2.51693361 | 0.585 | 0.059 | 0 | Endothelial | ENG |
| 2.49692911 | 0.506 | 0.038 | 0 | Endothelial | TMTC2 |
| 2.48158605 | 0.634 | 0.039 | 0 | Endothelial | FLNB |
| 4.44942402 | 0.915 | 0.01 | 0 | Smooth Muscle | CARMN |
| 4.33505611 | 0.95 | 0.073 | 0 | Smooth Muscle | CACNA1C |
| 4.25417169 | 0.817 | 0.039 | 0 | Smooth Muscle | GUCY1A2 |
| 4.18354928 | 0.789 | 0.059 | 0 | Smooth Muscle | EGFLAM |
| 4.00595646 | 0.962 | 0.184 | 0 | Smooth Muscle | DLC1 |
| 3.97619613 | 0.823 | 0.143 | 0 | Smooth Muscle | FRMD3 |
| 3.96928847 | 0.882 | 0.145 | 0 | Smooth Muscle | EPS8 |
| 3.61232024 | 0.863 | 0.063 | 0 | Smooth Muscle | PDGFRB |
| 3.60854355 | 0.924 | 0.147 | 0 | Smooth Muscle | NR2F2-AS1 |
| 3.57715006 | 0.74 | 0.172 | 0 | Smooth Muscle | PDZD2 |
| 3.48463003 | 0.76 | 0.099 | 0 | Smooth Muscle | AC012409.2 |
| 3.46687074 | 0.736 | 0.026 | 0 | Smooth Muscle | MYO1B |
| 3.44600111 | 0.836 | 0.089 | 0 | Smooth Muscle | RBPMS |
| 3.43181672 | 0.453 | 0.018 | 0 | Smooth Muscle | RGS6 |
| 3.42538949 | 0.664 | 0.019 | 0 | Smooth Muscle | RGS5 |
| 3.35267237 | 0.862 | 0.065 | 0 | Smooth Muscle | CLMN |
| 3.27194522 | 0.866 | 0.133 | 0 | Smooth Muscle | LHFPL6 |
| 3.22951613 | 0.963 | 0.283 | 0 | Smooth Muscle | CALD1 |
| 3.22784396 | 0.832 | 0.081 | 0 | Smooth Muscle | IGFBP7 |
| 3.09571113 | 0.558 | 0.03 | 0 | Smooth Muscle | PDE1C |
| 3.07129739 | 0.399 | 0.046 | 0 | Smooth Muscle | KCNAB1 |
| 3.06845726 | 0.682 | 0.074 | 0 | Smooth Muscle | PDE3A |
| 2.95817767 | 0.673 | 0.024 | 0 | Smooth Muscle | GUCY1A1 |
| 2.89523852 | 0.92 | 0.18 | 0 | Smooth Muscle | SOX5 |
| 2.84607813 | 0.551 | 0.024 | 0 | Smooth Muscle | ACTA2 |
| 2.84212635 | 0.687 | 0.022 | 0 | Smooth Muscle | NOTCH3 |
| 2.80804615 | 0.608 | 0.046 | 0 | Smooth Muscle | RIPOR3 |
| 2.76021099 | 0.863 | 0.26 | 0 | Smooth Muscle | RASAL2 |
| 2.75261641 | 0.597 | 0.062 | 0 | Smooth Muscle | CACNB2 |
| 2.74139594 | 0.502 | 0.024 | 0 | Smooth Muscle | AL356258.1 |
| 2.72717823 | 0.825 | 0.26 | 0 | Smooth Muscle | ADAMTS9-AS2 |
| 2.72356067 | 0.632 | 0.005 | 0 | Smooth Muscle | MRVI1 |
| 2.69416206 | 0.712 | 0.054 | 0 | Smooth Muscle | SPECC1 |
| 2.69086646 | 0.47 | 0.009 | 0 | Smooth Muscle | SLC16A12 |
| 2.66090378 | 0.549 | 0.001 | 0 | Smooth Muscle | FHL5 |
| 2.65191605 | 0.261 | 0.035 | 1.34E-231 | Smooth Muscle | MYH11 |
| 2.6278076 | 0.574 | 0.009 | 0 | Smooth Muscle | NFASC |
| 2.54897913 | 0.575 | 0.061 | 0 | Smooth Muscle | ADGRB3 |
| 2.53723162 | 0.399 | 0.007 | 0 | Smooth Muscle | AL499616.1 |
| 2.51667432 | 0.755 | 0.138 | 0 | Smooth Muscle | SPARCL1 |
| 2.49797483 | 0.562 | 0.068 | 0 | Smooth Muscle | ADAMTS12 |
| 2.48569963 | 0.413 | 0.058 | 0 | Smooth Muscle | SORBS2 |
| 2.46604675 | 0.729 | 0.243 | 0 | Smooth Muscle | SH3RF1 |
| 2.46517177 | 0.732 | 0.153 | 0 | Smooth Muscle | PLCB4 |
| 2.43854653 | 0.811 | 0.193 | 0 | Smooth Muscle | MAML2 |
| 2.43746213 | 0.853 | 0.371 | 0 | Smooth Muscle | INPP4B |
| 2.43373641 | 0.625 | 0.058 | 0 | Smooth Muscle | BMP5 |
| 2.43200493 | 0.537 | 0.035 | 0 | Smooth Muscle | COL5A3 |
| 2.42687747 | 0.421 | 0.085 | 1.47E-236 | Smooth Muscle | SLIT3 |
| 2.42023881 | 0.483 | 0.028 | 0 | Smooth Muscle | SEMA5A |
| 4.58539783 | 0.944 | 0.062 | 0 | Immune-B/T/NK 1 | ARHGAP15 |
| 4.39345782 | 0.911 | 0.014 | 0 | Immune-B/T/NK 1 | SKAP1 |
| 3.94825478 | 0.881 | 0.039 | 0 | Immune-B/T/NK 1 | PTPRC |
| 3.88885682 | 0.695 | 0.002 | 0 | Immune-B/T/NK 1 | THEMIS |
| 3.60609461 | 0.814 | 0.025 | 0 | Immune-B/T/NK 1 | IKZF1 |
| 3.49502095 | 0.835 | 0.156 | 0 | Immune-B/T/NK 1 | RIPOR2 |
| 3.44924432 | 0.45 | 0.031 | 0 | Immune-B/T/NK 1 | C15orf53 |
| 3.43060692 | 0.916 | 0.172 | 0 | Immune-B/T/NK 1 | ANKRD44 |
| 3.40560396 | 0.82 | 0.068 | 0 | Immune-B/T/NK 1 | PARP8 |
| 3.37477488 | 0.723 | 0.006 | 0 | Immune-B/T/NK 1 | SLFN12L |
| 3.31602016 | 0.506 | 0.012 | 0 | Immune-B/T/NK 1 | TOX |
| 3.2557841 | 0.694 | 0.001 | 0 | Immune-B/T/NK 1 | BCL11B |
| 3.25284525 | 0.659 | 0.01 | 0 | Immune-B/T/NK 1 | ITGA4 |
| 3.2003431 | 0.545 | 0.025 | 0 | Immune-B/T/NK 1 | PCAT1 |
| 3.18915584 | 0.751 | 0.03 | 0 | Immune-B/T/NK 1 | FYB1 |
| 3.14569407 | 0.511 | 0.005 | 0 | Immune-B/T/NK 1 | LINC01934 |
| 3.08792224 | 0.812 | 0.101 | 0 | Immune-B/T/NK 1 | CCND3 |
| 3.0680975 | 0.827 | 0.079 | 0 | Immune-B/T/NK 1 | DOCK8 |
| 3.05952162 | 0.596 | 0.01 | 0 | Immune-B/T/NK 1 | SAMD3 |
| 2.94460761 | 0.666 | 0.071 | 0 | Immune-B/T/NK 1 | ETS1 |
| 2.92154486 | 0.605 | 0.022 | 0 | Immune-B/T/NK 1 | CD247 |
| 2.91790315 | 0.728 | 0.084 | 0 | Immune-B/T/NK 1 | CHST11 |
| 2.90634838 | 0.728 | 0.105 | 0 | Immune-B/T/NK 1 | PIP4K2A |
| 2.8717821 | 0.593 | 0.004 | 0 | Immune-B/T/NK 1 | CARD11 |
| 2.86646911 | 0.684 | 0.031 | 0 | Immune-B/T/NK 1 | CD96 |
| 2.78930281 | 0.639 | 0.048 | 0 | Immune-B/T/NK 1 | PCED1B |
| 2.78048856 | 0.652 | 0.055 | 0 | Immune-B/T/NK 1 | TC2N |
| 2.73372717 | 0.613 | 0.039 | 0 | Immune-B/T/NK 1 | TNFAIP8 |
| 2.73230374 | 0.682 | 0.046 | 0 | Immune-B/T/NK 1 | APBB1IP |
| 2.72207928 | 0.58 | 0.007 | 0 | Immune-B/T/NK 1 | STAT4 |
| 2.67569812 | 0.555 | 0.001 | 0 | Immune-B/T/NK 1 | CD2 |
| 2.66425188 | 0.54 | 0.003 | 0 | Immune-B/T/NK 1 | ITK |
| 2.64950099 | 0.85 | 0.373 | 9.18E-268 | Immune-B/T/NK 1 | INPP4B |
| 2.64763958 | 0.481 | 0.005 | 0 | Immune-B/T/NK 1 | CAMK4 |
| 2.64199758 | 0.715 | 0.053 | 0 | Immune-B/T/NK 1 | DOCK2 |
| 2.61098457 | 0.42 | 0.001 | 0 | Immune-B/T/NK 1 | IL7R |
| 2.60772734 | 0.758 | 0.148 | 0 | Immune-B/T/NK 1 | DOCK10 |
| 2.57844833 | 0.73 | 0.055 | 0 | Immune-B/T/NK 1 | IQGAP2 |
| 2.55996946 | 0.705 | 0.113 | 0 | Immune-B/T/NK 1 | PITPNC1 |
| 2.51010371 | 0.507 | 0.005 | 0 | Immune-B/T/NK 1 | GRAP2 |
| 2.49834604 | 0.669 | 0.153 | 0 | Immune-B/T/NK 1 | KIAA1551 |
| 2.49470885 | 0.583 | 0.035 | 0 | Immune-B/T/NK 1 | MYO1F |
| 2.47176978 | 0.657 | 0.113 | 0 | Immune-B/T/NK 1 | CD44 |
| 2.45556258 | 0.519 | 0.01 | 0 | Immune-B/T/NK 1 | ITGAL |
| 2.4476903 | 0.779 | 0.179 | 0 | Immune-B/T/NK 1 | RUNX1 |
| 2.43986702 | 0.728 | 0.137 | 0 | Immune-B/T/NK 1 | FYN |
| 2.42326069 | 0.534 | 0.018 | 0 | Immune-B/T/NK 1 | CCDC88C |
| 2.42285417 | 0.652 | 0.163 | 1.03E-281 | Immune-B/T/NK 1 | MDFIC |
| 2.40896858 | 0.39 | 0.012 | 0 | Immune-B/T/NK 1 | PPP2R2B |
| 2.40432442 | 0.537 | 0.03 | 0 | Immune-B/T/NK 1 | PIK3R5 |
| 4.99242496 | 0.967 | 0.005 | 0 | Lymphatic Endothelial | PKHD1L1 |
| 4.52347936 | 0.979 | 0.087 | 0 | Lymphatic Endothelial | STOX2 |
| 4.2955808 | 0.869 | 0.003 | 0 | Lymphatic Endothelial | MMRN1 |
| 4.27070546 | 0.893 | 0.041 | 0 | Lymphatic Endothelial | ST6GALNAC3 |
| 4.24993682 | 0.8 | 0.009 | 0 | Lymphatic Endothelial | NRG3 |
| 4.23882344 | 0.943 | 0.1 | 0 | Lymphatic Endothelial | EFNA5 |
| 4.20372742 | 0.618 | 0.024 | 0 | Lymphatic Endothelial | AL357507.1 |
| 3.98515134 | 0.976 | 0.289 | 0 | Lymphatic Endothelial | PPFIBP1 |
| 3.88668032 | 0.734 | 0.009 | 0 | Lymphatic Endothelial | RELN |
| 3.5979962 | 0.731 | 0.008 | 0 | Lymphatic Endothelial | LINC02147 |
| 3.51750996 | 0.943 | 0.079 | 0 | Lymphatic Endothelial | PTPRE |
| 3.49966655 | 0.842 | 0.036 | 0 | Lymphatic Endothelial | NRP2 |
| 3.46421602 | 0.773 | 0.024 | 0 | Lymphatic Endothelial | PIEZO2 |
| 3.43037499 | 0.857 | 0.027 | 0 | Lymphatic Endothelial | SNTG2 |
| 3.42421188 | 0.91 | 0.118 | 0 | Lymphatic Endothelial | DOCK5 |
| 3.38201191 | 0.94 | 0.257 | 1.12E-283 | Lymphatic Endothelial | TFPI |
| 3.29114822 | 0.728 | 0.066 | 0 | Lymphatic Endothelial | GPM6A |
| 3.26185586 | 0.851 | 0.113 | 0 | Lymphatic Endothelial | PDE1A |
| 3.24128744 | 0.94 | 0.141 | 0 | Lymphatic Endothelial | TSHZ2 |
| 3.22183487 | 0.854 | 0.078 | 0 | Lymphatic Endothelial | RHOJ |
| 3.1230221 | 0.86 | 0.122 | 0 | Lymphatic Endothelial | ITGA9 |
| 3.11646232 | 0.785 | 0.064 | 0 | Lymphatic Endothelial | TSPAN5 |
| 3.09267968 | 0.693 | 0.044 | 0 | Lymphatic Endothelial | NTN1 |
| 3.08824138 | 0.648 | 0.012 | 0 | Lymphatic Endothelial | KLHL4 |
| 3.06246534 | 0.866 | 0.087 | 0 | Lymphatic Endothelial | LDB2 |
| 3.0451534 | 0.761 | 0.063 | 0 | Lymphatic Endothelial | CSGALNACT1 |
| 3.04203724 | 0.854 | 0.062 | 0 | Lymphatic Endothelial | VAV3 |
| 3.02695427 | 0.878 | 0.282 | 8.24E-217 | Lymphatic Endothelial | MPP7 |
| 3.01893033 | 0.579 | 0.015 | 0 | Lymphatic Endothelial | AC008691.1 |
| 3.01610349 | 0.761 | 0.017 | 0 | Lymphatic Endothelial | PARD6G |
| 2.95805784 | 0.824 | 0.067 | 0 | Lymphatic Endothelial | ELK3 |
| 2.94826253 | 0.863 | 0.28 | 2.44E-202 | Lymphatic Endothelial | PGM5 |
| 2.93784994 | 0.767 | 0.09 | 0 | Lymphatic Endothelial | PROX1 |
| 2.91424754 | 0.878 | 0.151 | 0 | Lymphatic Endothelial | KIAA1671 |
| 2.86462199 | 0.779 | 0.085 | 0 | Lymphatic Endothelial | HMCN1 |
| 2.85990005 | 0.815 | 0.176 | 4.75E-271 | Lymphatic Endothelial | SMAD1 |
| 2.85561125 | 0.916 | 0.434 | 1.06E-179 | Lymphatic Endothelial | KALRN |
| 2.85183049 | 0.642 | 0.01 | 0 | Lymphatic Endothelial | STAB2 |
| 2.82021757 | 0.436 | 0.016 | 0 | Lymphatic Endothelial | FAM155A |
| 2.81408621 | 0.967 | 0.484 | 5.51E-192 | Lymphatic Endothelial | MAGI1 |
| 2.81257851 | 0.752 | 0.07 | 0 | Lymphatic Endothelial | STK32B |
| 2.796392 | 0.904 | 0.242 | 3.71E-254 | Lymphatic Endothelial | ZNF521 |
| 2.77675979 | 0.857 | 0.102 | 0 | Lymphatic Endothelial | LAMA4 |
| 2.77077765 | 0.878 | 0.155 | 0 | Lymphatic Endothelial | NR2F2-AS1 |
| 2.74305139 | 0.579 | 0.023 | 0 | Lymphatic Endothelial | SYT1 |
| 2.73714741 | 0.681 | 0.029 | 0 | Lymphatic Endothelial | NALCN |
| 2.71539914 | 0.722 | 0.003 | 0 | Lymphatic Endothelial | FLT4 |
| 2.69439057 | 0.857 | 0.091 | 0 | Lymphatic Endothelial | COLEC12 |
| 2.6790771 | 0.597 | 0.019 | 0 | Lymphatic Endothelial | TLL1 |
| 2.63839586 | 0.657 | 0.016 | 0 | Lymphatic Endothelial | NR2F1-AS1 |
| 4.90560416 | 0.763 | 0.064 | 0 | Immune-B/T/NK 2 | NTM |
| 4.76039079 | 0.9 | 0.002 | 0 | Immune-B/T/NK 2 | CPA3 |
| 4.43444342 | 0.819 | 0.012 | 0 | Immune-B/T/NK 2 | HPGD |
| 4.40141192 | 0.884 | 0.002 | 0 | Immune-B/T/NK 2 | KIT |
| 4.06469981 | 0.855 | 0.069 | 0 | Immune-B/T/NK 2 | STX3 |
| 3.96471554 | 0.819 | 0.001 | 0 | Immune-B/T/NK 2 | HDC |
| 3.94671378 | 0.795 | 0.002 | 0 | Immune-B/T/NK 2 | MS4A2 |
| 3.84859005 | 0.743 | 0.01 | 0 | Immune-B/T/NK 2 | IL18R1 |
| 3.76377057 | 0.731 | 0.004 | 0 | Immune-B/T/NK 2 | RAB27B |
| 3.75729715 | 0.783 | 0.023 | 0 | Immune-B/T/NK 2 | HPGDS |
| 3.64674922 | 0.747 | 0.073 | 0 | Immune-B/T/NK 2 | MEIS2 |
| 3.59711351 | 0.88 | 0.225 | 3.15E-205 | Immune-B/T/NK 2 | BMP2K |
| 3.51941131 | 0.691 | 0.008 | 0 | Immune-B/T/NK 2 | CDK15 |
| 3.49745174 | 0.751 | 0.023 | 0 | Immune-B/T/NK 2 | ALOX5 |
| 3.46577517 | 0.502 | 0.01 | 0 | Immune-B/T/NK 2 | LINC02147 |
| 3.40495436 | 0.767 | 0.069 | 0 | Immune-B/T/NK 2 | BACE2 |
| 3.37527028 | 0.964 | 0.371 | 1.59E-175 | Immune-B/T/NK 2 | SLC24A3 |
| 3.32225361 | 0.703 | 0.01 | 0 | Immune-B/T/NK 2 | MCTP2 |
| 3.27922936 | 0.843 | 0.07 | 0 | Immune-B/T/NK 2 | ARHGAP15 |
| 3.27120623 | 0.703 | 0.062 | 0 | Immune-B/T/NK 2 | SGK1 |
| 3.25872722 | 0.635 | 0.013 | 0 | Immune-B/T/NK 2 | KIAA1549 |
| 3.24281482 | 0.888 | 0.187 | 1.83E-223 | Immune-B/T/NK 2 | ELMO1 |
| 3.20650938 | 0.695 | 0.051 | 0 | Immune-B/T/NK 2 | ST8SIA1 |
| 3.18993484 | 0.627 | 0.001 | 0 | Immune-B/T/NK 2 | TPSB2 |
| 3.17818409 | 0.643 | 0.023 | 0 | Immune-B/T/NK 2 | SKAP1 |
| 3.13274014 | 0.606 | 0.004 | 0 | Immune-B/T/NK 2 | GATA2 |
| 3.1164897 | 0.594 | 0.005 | 0 | Immune-B/T/NK 2 | RGS13 |
| 3.09355811 | 0.578 | 0.011 | 0 | Immune-B/T/NK 2 | SLC18A2 |
| 3.08856867 | 0.643 | 0.01 | 0 | Immune-B/T/NK 2 | VWA5A |
| 3.07598124 | 0.783 | 0.252 | 1.48E-136 | Immune-B/T/NK 2 | SYTL3 |
| 3.05434935 | 0.699 | 0.169 | 1.19E-136 | Immune-B/T/NK 2 | LRP1B |
| 3.01661037 | 0.727 | 0.089 | 4.67E-290 | Immune-B/T/NK 2 | CHST11 |
| 3.00787946 | 0.811 | 0.179 | 6.03E-190 | Immune-B/T/NK 2 | ANKRD44 |
| 2.96714212 | 0.835 | 0.327 | 6.97E-121 | Immune-B/T/NK 2 | FER |
| 2.91574607 | 0.602 | 0.053 | 0 | Immune-B/T/NK 2 | SLCO2B1 |
| 2.89459565 | 0.586 | 0.016 | 0 | Immune-B/T/NK 2 | TOX |
| 2.85602336 | 0.59 | 0.048 | 0 | Immune-B/T/NK 2 | CPM |
| 2.83349071 | 0.578 | 0.002 | 0 | Immune-B/T/NK 2 | RHEX |
| 2.8236518 | 0.691 | 0.118 | 2.55E-194 | Immune-B/T/NK 2 | CD44 |
| 2.8138847 | 0.582 | 0.04 | 0 | Immune-B/T/NK 2 | PPM1H |
| 2.79927469 | 0.59 | 0.03 | 0 | Immune-B/T/NK 2 | KCNQ1 |
| 2.79675862 | 0.663 | 0.089 | 2.07E-243 | Immune-B/T/NK 2 | ACER3 |
| 2.79330925 | 0.92 | 0.199 | 1.17E-208 | Immune-B/T/NK 2 | LRMDA |
| 2.78568836 | 0.57 | 0.033 | 0 | Immune-B/T/NK 2 | PRKCB |
| 2.77057572 | 0.418 | 0.002 | 0 | Immune-B/T/NK 2 | AC092979.1 |
| 2.74207108 | 0.53 | 0.092 | 2.37E-140 | Immune-B/T/NK 2 | LMO4 |
| 2.72313604 | 0.462 | 0.009 | 0 | Immune-B/T/NK 2 | ENPP3 |
| 2.68334349 | 0.59 | 0.08 | 4.13E-203 | Immune-B/T/NK 2 | STXBP6 |
| 2.68030745 | 0.582 | 0.047 | 0 | Immune-B/T/NK 2 | SYTL2 |
| 2.67541307 | 0.49 | 0.002 | 0 | Immune-B/T/NK 2 | P2RX1 |
| 5.36782924 | 0.976 | 0.106 | 0 | Adipocytes | GPAM |
| 5.11736617 | 0.964 | 0.037 | 0 | Adipocytes | PDE3B |
| 4.88301091 | 0.976 | 0.047 | 0 | Adipocytes | PPARG |
| 3.92215437 | 0.982 | 0.009 | 0 | Adipocytes | PLIN1 |
| 3.91954907 | 0.911 | 0.109 | 3.42E-271 | Adipocytes | GRK3 |
| 3.80091171 | 0.976 | 0.104 | 0 | Adipocytes | LAMA4 |
| 3.69602352 | 0.94 | 0.046 | 0 | Adipocytes | MGST1 |
| 3.5616431 | 0.464 | 0.004 | 0 | Adipocytes | SCD |
| 3.54280225 | 0.94 | 0.126 | 2.87E-268 | Adipocytes | PLIN4 |
| 3.49705007 | 0.982 | 0.19 | 2.06E-200 | Adipocytes | SOX5 |
| 3.4906178 | 0.923 | 0.032 | 0 | Adipocytes | TMEM132C |
| 3.47560177 | 0.869 | 0.125 | 1.48E-211 | Adipocytes | LPL |
| 3.407356 | 0.988 | 0.201 | 3.20E-188 | Adipocytes | EBF1 |
| 3.36884315 | 0.839 | 0.058 | 0 | Adipocytes | LIPE-AS1 |
| 3.32632377 | 0.994 | 0.558 | 4.05E-108 | Adipocytes | ACACB |
| 3.1786584 | 0.893 | 0.052 | 0 | Adipocytes | APBB1IP |
| 3.15717126 | 0.631 | 0.009 | 0 | Adipocytes | FASN |
| 3.11706215 | 0.833 | 0.039 | 0 | Adipocytes | PLXNA4 |
| 3.11107694 | 0.857 | 0.002 | 0 | Adipocytes | ADIPOQ |
| 3.10847807 | 0.839 | 0.009 | 0 | Adipocytes | LINC02237 |
| 3.04258436 | 0.905 | 0.355 | 1.10E-86 | Adipocytes | ACSL1 |
| 2.9869413 | 0.768 | 0.042 | 0 | Adipocytes | LINC00598 |
| 2.95486347 | 0.774 | 0.007 | 0 | Adipocytes | PTGER3 |
| 2.88887027 | 0.881 | 0.048 | 0 | Adipocytes | CPM |
| 2.86773012 | 0.881 | 0.067 | 0 | Adipocytes | ADH1B |
| 2.84045137 | 0.952 | 0.318 | 5.78E-105 | Adipocytes | EHBP1 |
| 2.83304415 | 0.792 | 0.014 | 0 | Adipocytes | PRKAR2B |
| 2.82068605 | 0.905 | 0.134 | 3.54E-212 | Adipocytes | SLC16A7 |
| 2.80565482 | 0.929 | 0.18 | 6.67E-174 | Adipocytes | MLXIPL |
| 2.74118191 | 0.893 | 0.103 | 1.06E-275 | Adipocytes | PNPLA2 |
| 2.69133986 | 0.702 | 0.024 | 0 | Adipocytes | AC002066.1 |
| 2.69079183 | 0.774 | 0.051 | 0 | Adipocytes | MAOA |
| 2.67890171 | 0.887 | 0.04 | 0 | Adipocytes | PECR |
| 2.67344819 | 0.935 | 0.164 | 8.32E-187 | Adipocytes | DIRC3 |
| 2.66557903 | 0.708 | 0.01 | 0 | Adipocytes | FABP4 |
| 2.64797424 | 0.929 | 0.158 | 1.40E-205 | Adipocytes | AQP7 |
| 2.62279373 | 0.952 | 0.394 | 2.96E-99 | Adipocytes | SIK2 |
| 2.61782189 | 0.679 | 0.046 | 0 | Adipocytes | ELOVL5 |
| 2.6026313 | 0.887 | 0.173 | 5.74E-161 | Adipocytes | MAST4 |
| 2.59754817 | 0.958 | 0.476 | 1.06E-97 | Adipocytes | TNS1 |
| 2.5681249 | 0.851 | 0.022 | 0 | Adipocytes | CARMN |
| 2.56075735 | 0.863 | 0.106 | 5.28E-215 | Adipocytes | GPC6 |
| 2.54925524 | 0.899 | 0.231 | 2.04E-122 | Adipocytes | VKORC1L1 |
| 2.53267729 | 0.786 | 0.002 | 0 | Adipocytes | GYG2 |
| 2.52606078 | 0.685 | 0.015 | 0 | Adipocytes | ADRA1A |
| 2.51685927 | 0.69 | 0.104 | 2.88E-142 | Adipocytes | CLSTN2 |
| 2.51179905 | 0.792 | 0.052 | 0 | Adipocytes | TRHDE |
| 2.50294951 | 0.732 | 0.023 | 0 | Adipocytes | LIPE |
| 2.45768721 | 0.554 | 0.035 | 7.56E-279 | Adipocytes | AC104574.2 |
| 2.45121823 | 0.893 | 0.368 | 1.11E-74 | Adipocytes | EEPD1 |
